# Fluctuating environments maintain genetic diversity through neutral fitness effects and balancing selection

**DOI:** 10.1101/2021.02.23.432553

**Authors:** Farah Abdul-Rahman, Daniel Tranchina, David Gresham

## Abstract

Genetic variation is the raw material upon which selection acts. The majority of environmental conditions change over time and therefore may result in variable selective effects. How temporally fluctuating environments impact the distribution of fitness effects and in turn population diversity is an unresolved question in evolutionary biology. Here, we employed continuous culturing using chemostats to establish environments that switch periodically between different nutrient limitations and compared the dynamics of selection to static conditions. We used the pooled *Saccharomyces cerevisiae* haploid gene deletion collection as a synthetic model for populations comprising thousands of unique genotypes. Using barcode sequencing (barseq), we find that static environments are uniquely characterized by a small number of high fitness genotypes that rapidly dominate the population leading to dramatic decreases in genetic diversity. By contrast, fluctuating environments are enriched in genotypes with neutral fitness effects and an absence of extreme fitness genotypes contributing to the maintenance of genetic diversity. We also identified a unique class of genotypes whose frequencies oscillate sinusoidally with a period matching the environmental fluctuation. Oscillatory behavior corresponds to large differences in short term fitness that are not observed across long timescales pointing to the importance of balancing selection in maintaining genetic diversity in fluctuating environments. Our results are consistent with a high degree of environmental specificity in the distribution of fitness effects and the combined effects of reduced and balancing selection in maintaining genetic diversity in the presence of variable selection.

## Introduction

The dynamics of adaptive evolution in genetically heterogeneous populations depend on the strength of selection and the distribution of fitness effects among genotypes (Bell 2008). How selective environmental conditions and genetic variation contribute to evolutionary dynamics is one of the central questions in evolutionary biology. In genetically heterogeneous populations the fitness of different genotypes varies and selection acts to increase the relative abundance of advantageous genotypes. In the simplest scenario, comprising a single fitness component (i.e. a single selective force) and single variant differences between genotypes, the distribution of fitness effects can reliably predict the dynamics of adaptive evolution. However, the impact of variable selective conditions, that result from fluctuations in the environment, on the distribution of fitness effects and the corresponding impact on genetic diversity is poorly understood.

In natural and engineered environments, organisms frequently encounter fluctuating selection in the form of physical or biotic factors (Bell 2010; Nguyen et al. 2020). Fluctuations in environmental conditions may occur with a regular period, such as diurnal fluctuations or aperiodically, such as with the random temperature variations that occur throughout the day. Periodic environmental fluctuations comprise an enormous range of timescales and patterns ranging from hours, as is the case with the gut microbiome (Schlomann and Parthasarathy 2019), to millenia, such as the timing between glacial periods. The prevalence of periodic fluctuations at different time scales in diverse environments suggests that our understanding of how evolution has shaped extant organisms and our ability to predict the future outcomes of adaptation requires understanding how organisms respond to environmental change.

The molecular basis of adaptation depends on the selective regime of the fluctuating environment. Regularly repeating and predictable fluctuations have been shown to select for anticipatory strategies such as a memory-dependent response (Razinkov et al. 2013; Lambert and Kussell 2014) or programmed oscillatory behavior (Johnson and Golden 2006). Conversely, fluctuations that occur at random intervals may favor strategies that don’t rely on forecasting future environmental conditions, such as sense-and-response (Uschner and Klipp 2014) or bet-hedging strategies (Olofsson et al. 2009; Childs et al. 2010). The frequency of environmental fluctuation with respect to generation time is also a key determinant of adaptive strategies; if the fluctuations are extremely rapid with respect to generation time, an organism may sense a time averaged condition, whereas extremely slow oscillations with respect to generation time may result in organisms experiencing effectively static conditions (Cvijović et al. 2015; Lin and Kussell 2016). Moderate fluctuations that fall between the two extremes and with periods less than the generation time likely apply selective pressure on regulatory pathways requiring an organism to respond to environmental change on the individual level.

A variety of theoretical expectations underlie the selective dynamics of genotypes in fluctuating environments. Balancing selection, generally defined as any type of selection that maintains polymorphism in a population, can explain the maintenance of genetic diversity in temporally varying environments. For example, fluctuations with periods over multiple generations can select for the coexistence of genotypes specialized to each of the two environments (Bergland et al. 2014; New et al. 2014) an ecological phenomenon known as the “Temporal Storage Effect” (Chesson 1994; Tan et al. 2017; Letten et al. 2018). In the extreme case, antagonistic pleiotropy, in which an allele that is beneficial in one condition is deleterious in another, can manifest between the different phases of a periodically fluctuating environment. By contrast, a neutralist view aims to explain the maintenance of variation in fluctuating environments by a combination of other processes such as the continual generation of mutation in a population and the introduction of variation through migration (Bertram and Masel 2019)Hedrik et al. 1976; (Bertram and Masel 2019). Theoretical analyses of fluctuating environments have suggested that the efficiency of selection can be reduced in variable environments (Cvijović et al. 2015; Cvijovic et al.). It has also been proposed that varying environments themselves can trigger increased mutation rate and thereby increase population diversity (Nelson and Masel 2018), or act as catalysts for evolution through more efficient phenotypic exploration (Kashtan et al. 2007).

Empirical approaches to studying selection in fluctuating environments present several challenges. In natural environments, experimental variables are difficult to control. Experimental evolution in a lab setting provides a potentially powerful approach and as such a number of studies have investigated the effect of fluctuating environments on adaptive evolution using experimental evolution of microbes (Cooper and Lenski 2010). In general, experimental microbial evolution studies have focused on genetically homogeneous populations and the effect of *de novo* mutation. However, a small number of studies have made use of genetically heterogeneous populations to address effects of environmental fluctuations (Salignon et al. 2018). To the best of our knowledge, all studies of microbial evolution reported to date have used batch culture, which requires serial passaging and population bottlenecking, adding additional variables to the study design. The precise control of selection in batch culture is also challenging because nutrient concentration changes continuously with population expansion even when the culture medium is unchanged (Li et al. 2018).

To study the effect of environmental fluctuations on the dynamics of adaptive evolution, we used the chemostat to establish different selective regimes and study their effect on population genetic diversity and the distribution of fitness effects in *Saccharomyces cerevisiae*. We used synthetic populations consisting of the pooled nonessential haploid gene deletion library (~4,000 unique genotypes) and quantified population composition using barcode sequencing (barseq). We characterized fluctuating environments in the chemostat using both experimental studies and mathematical modeling. We find that environments, in which nutrient concentration fluctuates, maintain greater genetic diversity than static environments. Increased genetic diversity in fluctuating environments results from the absence of a small number of highly fit and specialized genotypes that rapidly dominate populations evolving in static conditions and an enrichment in fluctuating environments of genotypes with neutral fitness effects. Many genotypes show non-linear and non-monotonic responses (log abundance versus time) to both static and fluctuating selection, but fluctuating environments uniquely select for a class of genotypes with oscillatory growth behavior. Oscillatory behavior contributes to large short term fitness effects that are not observed over the long term. Our study highlights the importance of environmental variability in evolutionary dynamics and provides a framework for modeling evolutionary scenarios that better reflect natural environments.

## Results

The empirical study of adaptive evolution requires consideration of both the selective conditions and the heritable variation in a population. In this study, we combined continuous culturing using chemostats and the *Saccharomyces cerevisiae* haploid non essential gene deletion collection to study the effect of temporally fluctuating selection on standing genetic variation.

### Modeling nutritional fluctuations in chemostats

Chemostats operate through continuous addition of fresh medium and removal of culture at the same rate (**Figure 1A**). We focused on two static conditions, carbon limitation (low carbon, high nitrogen) using glucose as the sole carbon source and nitrogen limitation (high carbon, low nitrogen) using ammonium sulfate as the sole nitrogen source. We constructed a periodically fluctuating condition in which the medium was switched between the two nutrient limiting conditions (**Figure 1B**). In this switch condition, the feed media alternates between the carbon limiting and nitrogen limiting media every 30 hours and the change in the feed media is instantaneous. We used the standard chemostat model (Monod 1950; Novick and Szilard 1950) to predict changes in nutrient concentration for single-nutrient limitation. We extended this model (**methods**) to incorporate temporal fluctuations in nutrient concentration and constrained cellular growth with a parameter for a second nutrient (Boer et al. 2010) to account for both changes in carbon and nitrogen concentrations.

**Figure 1.**
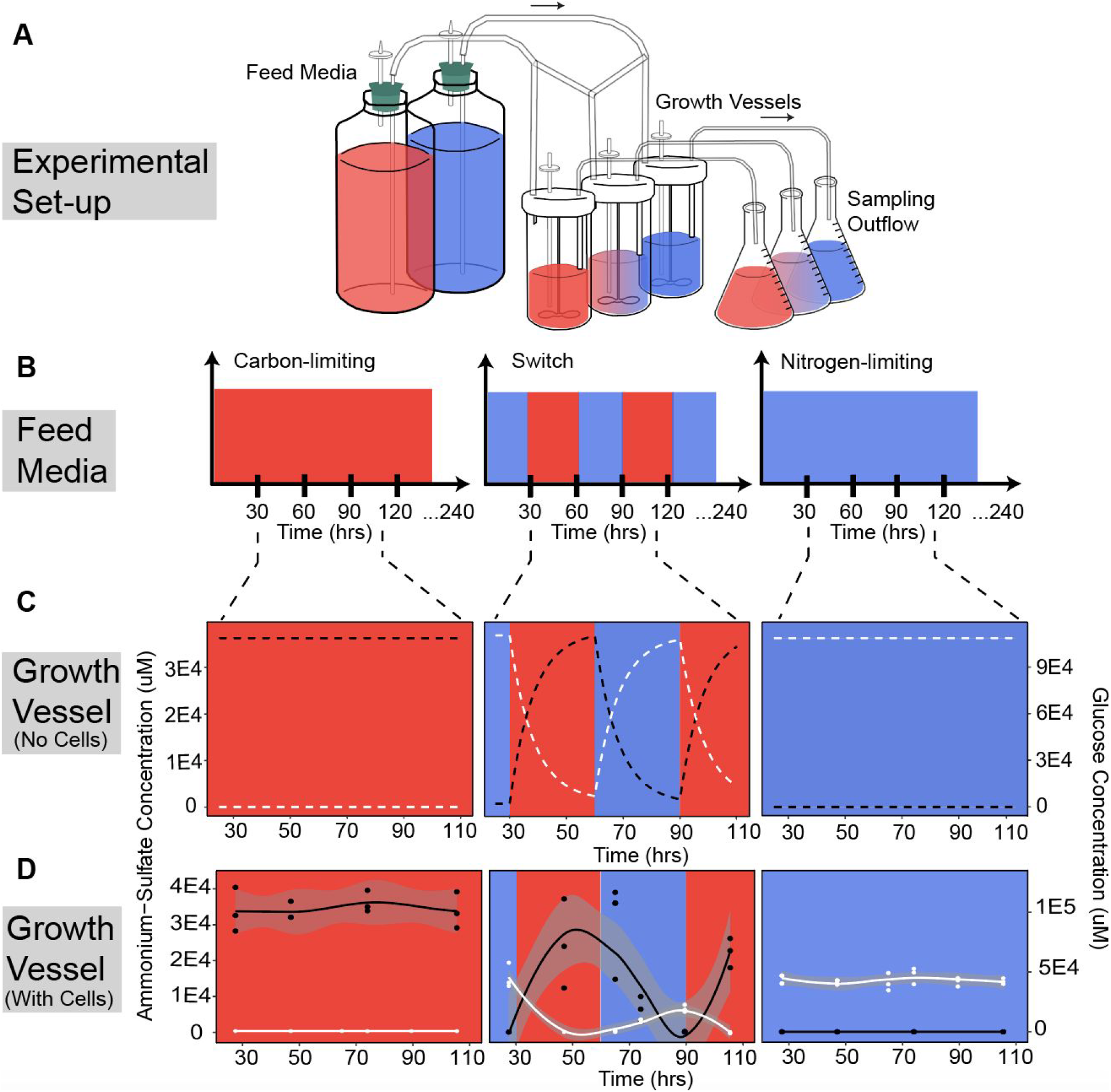
Periodically fluctuating environments in the chemostat. (**A**) We used chemostat cultures to study evolutionary dynamics in static and fluctuating conditions. (**B**) Populations were cultured in either carbon-limited media, nitrogen-limited media, or media that switched between the two nutrient limiting conditions every 30 hours (i.e. a period of 60 hours). All selections were maintained for a total of 240 hours. (**C**) An ordinary differential equation model was used to determine the expected concentrations of glucose (white), the sole carbon source, and ammonium sulfate (black), the sole nitrogen source, in the culture vessels in the absence of cellular consumption. (**D**) We experimentally measured glucose (white) and ammonium sulfate (black) concentrations in each of the culture vessels to determine the contribution of cellular consumption to environmental nutrient concentrations.

We first modelled nutrient concentrations in the chemostat in the absence of cells to study the effect of dilution alone. Whereas a single limiting nutrient results in a constant nutrient concentration, switching the media results in oscillations in nutrient concentration in the growth vessel that follow first-order (exponential) kinetics despite instantaneous switches in the feed media (**Figure 1C**). We then inoculated chemostats with a single wildtype genotype and measured ammonium-sulfate and glucose concentrations in each of the culture vessels during steady-state cellular growth to determine the effect of cellular consumption on nutrient concentration in the chemostat (**Figure 1D**). As expected, in all cases cellular consumption results in reduced nutrient concentrations in the chemostat. In the switch condition we find that the ammonium sulfate concentration oscillates between maximal and minimal levels that are equivalent to those observed in the two static conditions. By contrast, the maximal glucose concentration in the switch condition is reduced compared to glucose levels observed in static nitrogen limitation once the oscillations commence. This suggests that cells that have been previously exposed to growth-limiting levels of glucose exhibit increased glucose consumption in glucose rich conditions compared with cells that have not previously experienced growth-limiting glucose concentrations. This memory-like response may be due to the sustained expression of high affinity glucose transporters, induced by exposure to growth limiting glucose concentrations in the previous phase, persisting into the ammonium-sulfate limited phase (Buziol et al. 2008; Rintala et al. 2008).

We also considered an additional type of fluctuating environmental condition that differs in frequency and form. This fluctuation consisted of growth in steady-state ammonium-sulfate limiting conditions to which a bolus of 80uM nitrogen, in the form of either ammonium-sulfate or glutamine, was provided every three hours. This minor environmental perturbation, which we refer to as a “pulse” has previously been employed in studying transcriptional responses to environmental perturbation (Ronen and Botstein 2006; Airoldi et al. 2016). Prior mathematical modeling of chemostat pulses indicates that they result in a transient perturbation of the environment that rapidly returns to the steady-state condition (Airoldi et al. 2016).

### Selection on heterogeneous populations in a chemostat

We sought to quantify the dynamics of thousands of genotypes in static and fluctuating environments using chemostats. Classical chemostat theory holds that through the process of continuous growth and dilution, a chemostat attains a steady state in which the growing population size is constant and the concentration of the growth rate limiting nutrient is constant (Monod 1950; Novick and Szilard 1950; Kubitschek 1970). At steady-state, the population mean exponential growth-rate constant (*λ*) is equal to the chemostat dilution rate (*β*). However, competition for the limiting resource between the thousands of genotypes present in our experiments means that growth rates differ between genotypes. In our experiment, the growth rate of an individual genotype *i*, *λ_i_*, is defined as the fitness of genotype *i*. Fitness differences across genotypes result in corresponding changes in population proportions over time. Intuitively, one might think that the changing proportions of genotypes would preclude constancy of the population growth rate. How can the constancy of population growth rate in the chemostat (*λ* = *β*) be reconciled with the presence of thousands of distinct genotypes with different fitness effects?

To address this question we modelled the growth of 4,000 genotypes in a nutrient-limited chemostat based on a straightforward extension of the two-genotype model of competitive growth in a chemostat from (Dean 2005) (**supplemental methods**). As with Dean’s two-genotype model, we find that the total population size and concentration of the limiting nutrient do in fact change as selection acts on the different genotypes. However, these changes are negligible after an initial transient period. We find that in the case of a static environmental selection in the chemostat, the genotype proportions change until a steady state is ultimately achieved in which only a single growth rate constant remains in the chemostat. In this new steady-state condition the population size is increased and the growth limiting nutrient concentration is decreased relative to the initial conditions (**supplemental figure 1**). As the preceding period during which selection takes place is not a true steady-state, we refer to the selection during this time period as occurring in quasi steady-state conditions.

We also applied the Price equation in the continuous form (Day et al. 2020) to this scenario and found that the population growth rate cannot be exactly constant until the overall steady state condition above is achieved (**supplemental figure 1**). Examination of the Price equation shows that evolution of the population growth rate is driven by the variance of the growth rate and the rates of change of genotype fitness (**supplemental methods**).

### Fluctuating environments maintain greater genetic diversity

The distribution of fitness effects (DFE) quantitatively describes the proportion of variants in a population that are advantageous, neutral or deleterious, compared to the arithmetic mean fitness of the population. The shape of the DFE is influenced by several factors including the type of species, population size, and genome size (Eyre-Walker and Keightley 2007). Whereas both theoretical (Connallon and Clark 2015) and experimental (Cooper and Lenski 2010; Hietpas et al. 2013; Blundell et al. 2017) studies have investigated the effect of a variety of environments on the DFE, the effect of temporal environmental variation on the DFE remains largely unknown. Moreover, the consequences of variable selection on the maintenance of genetic diversity is poorly understood.

To address the effect of variable selection on the DFE and genetic diversity we used an isogenic single-gene deletion library to compare selection in static and fluctuating environments. The presence of unique molecular barcodes enables identification of ~4000 genotypes using quantitative DNA barcode sequencing (barseq) (Delneri 2010). We used the haploid gene deletion collection and barseq to quantify population diversity and genotype fitness over approximately forty generations (240 hours) of sustained selection (**Figure 2A**). By replicating selection experiments and limiting their duration, our approach minimizes the potential confounding effect of *de novo* mutations. Assuming a rate of 2.7 × 10^−3^ mutations/genome/replication (Drake et al. 1998) we would expect 0.108 mutations/ genome over 40 generations. Consistent with this expectation, after filtering sequencing libraries (**Supplemental figures 2A** **and** **2B**, **Supplemental table 1)**, replicate populations showed high within-condition correlation indicating that *de novo* mutations did not play a significant role in selection dynamics. A small number of replicate experiments with low correlation were excluded from further analysis (**Supplemental figure 2C**). Following quality control, our study comprised 278 barseq libraries.

**Figure 2.**
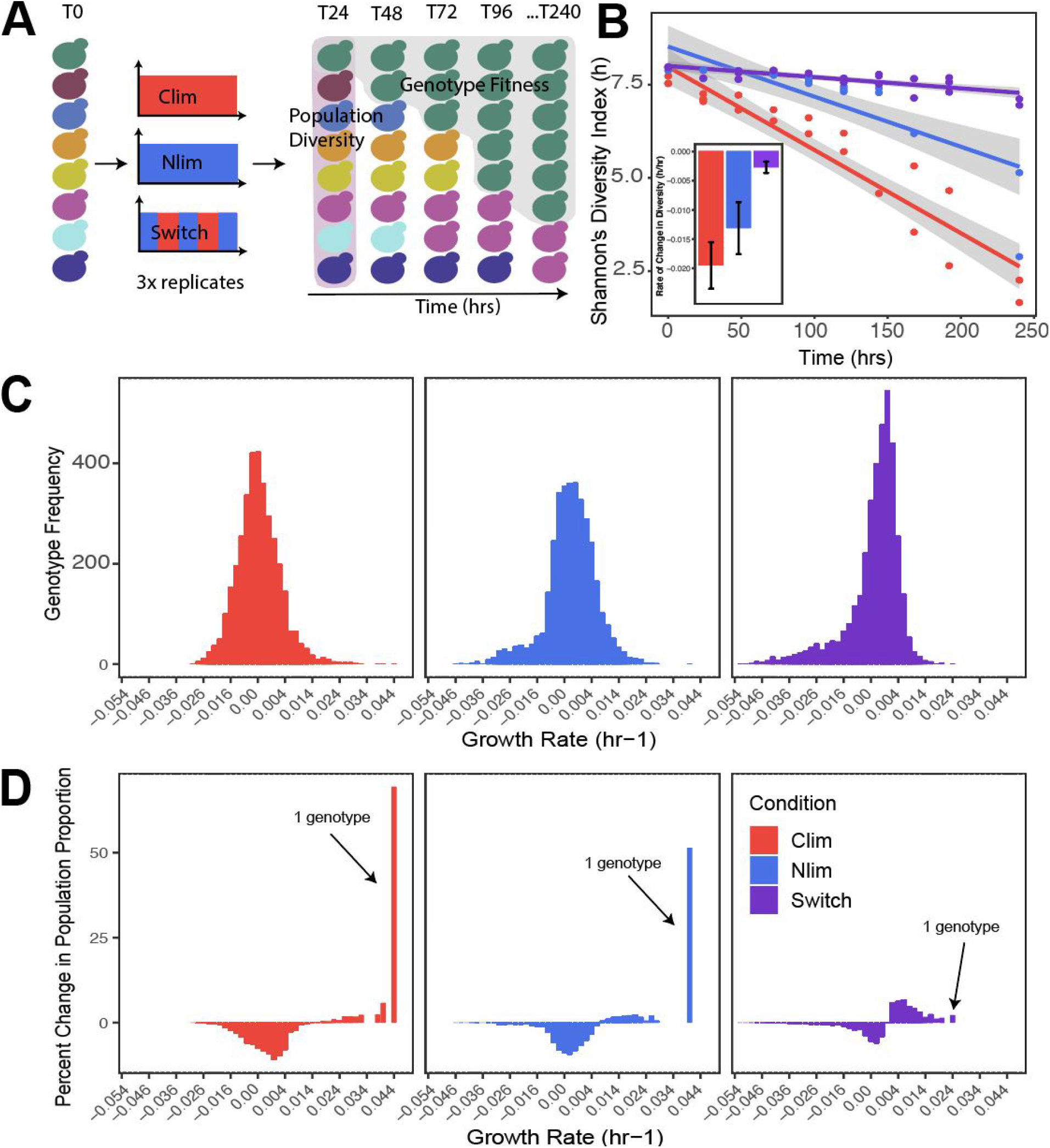
Fluctuating selective conditions maintain greater genetic diversity than static selective conditions. A single-gene deletion library containing ~4000 distinct gene knockout mutants was grown for 240 hours (approximately 40 generations) in static carbon-limitation, static nitrogen-limitation, and switching conditions in biological triplicate. Populations were sampled every 24 hours for a total of 10 timepoints. Barseq was used to estimate relative genotype abundance at each time point (**methods**). (**A**) Population diversity and genotype fitness were computed based on changes in mutant abundance across time (see **supplemental methods**). (**B**) The changes in Shannon’s diversity index show a clear reduction in population diversity over time in static conditions, but not in a fluctuating environment. (**C**) The distribution of growth rates, relative to the arithmetic mean over all genotypes, for ~4000 genotypes in each condition estimated over the complete 240 hours of growth and (**D**) the change in the population proportion within each growth rate bin between t = 0 and t = 240 hours.

To test the effect of environmental variability on population diversity we estimated the normalized abundance of each genotype at each time point in each condition (**supplemental methods**). We quantified the temporal change per unit time (in hours) rather than per generation to enable direct comparison between conditions as population growth rates in fluctuating chemostats are not necessarily determined by the dilution rate as they are in static conditions. We quantified population diversity using Shannon’s diversity index, which takes into consideration the richness of genotypes and the evenness of their abundances. We find that the static carbon-limiting and nitrogen-limiting conditions display rapid declines in diversity in comparison to the switch condition (**Figure 2B**). To test the generality of this conclusion we applied the same analysis to the two pulse conditions. In the presence of pulsed perturbations, populations also maintained greater genetic diversity across time suggesting that this feature may be generalizable to different frequencies and forms of environmental fluctuation (**supplemental figure 3**). We found that diversity estimates are not significantly affected by barcode library size (pearson r = 0.106, p-value = 0.218) (**supplemental figure 4**) excluding confounding effects of library size on diversity metrics. In addition, population richness does not appear to undergo large changes over time in any selection regime suggesting that differences in diversity are largely driven by changes in evenness (**supplemental figure 5**). All selections resulted in some degree of genotype extinction, defined by the absence of a genotype in a one or more terminal time points. We did not identify a common set of extinct genotypes across all conditions (**supplemental figure 6)**.

To test if differences in the rate of change in genetic diversity are associated with differences in fitness effects we computed the DFE for each condition. To quantify fitness over a given time interval (t1, t2) we use the temporal mean growth rate per cell minus the arithmetic mean over all genotypes. This is given by the difference between the log of normalized abundance at the two time points divided by the time difference (**supplemental methods**). By using the chemostat, the population exponential growth rate constant is set at 0.12 hr^−1^, which is equal to the population mean growth rate over all genotypes to the extent that the total number of cells in the chemostat remains constant (**supplemental methods**). We calculated average genotype fitness using the first (t = 0 hours) and last (t = 240 hours) time point. The DFEs in each condition have similar distributions characterized by a unimodal distribution centered around neutral relative fitness (**Figure 2C**). The DFE in all three conditions comprises primarily neutral genotypes with tails reflecting deleterious and beneficial genotypes relative to the mean population fitness. This property also holds for pulse fluctuations (**supplemental figure 7**). Whereas measures of dispersion for each DFE are similar between conditions, contrary to previous predictions (Connallon and Clark 2015), static conditions are distinguished by the presence of individual genotypes with extreme fitness effects (**supplemental table 2**). Thus, the distinguishing feature of the DFE, calculated over the entire period of selection, in static populations is the occurrence of extreme high fitness genotypes that are not observed in fluctuating selections. This observation is consistent with theoretical analysis using the Price equation (**supplemental methods**)

The presence of a single particularly high fitness genotype results in a distinct population composition following 240 hours of selection. In both static selection conditions, a single highly fit genotype comprises over 50% of the population at this terminal timepoint (**Figure 2D**). By contrast, the maximal frequency of the highest fitness genotype is only 3% in the switch condition (**supplemental table 2)**. In pulse fluctuations, the increased frequency of a small number of genotypes in the populations is apparent; however, this effect is reduced compared with static conditions (**supplemental figure 7** and **supplemental table 2**). These results point to a clear causal relationship between the presence of a single high fitness genotype and a rapid reduction in genetic diversity in static environments in which a single dominant selective pressure acts.

To test the generality of our observations we analyzed the dataset of *Salingon et al.* (Salignon et al. 2018) who studied the single-gene deletion library in two types of fluctuating environments using serial batch cultures and bottlenecking. In one of the fluctuating conditions (temporal variation in methionine concentration) we observed the same trend as our study. However, in the case of environments that fluctuate in salt concentration we find the opposite trend (**supplemental figure 8**). In this case, it is possible that specific gene deletions (e.g. *CIN5Δ/Δ, YOR028WΔ/Δ, SRFI1Δ/Δ)* are uniquely able to respond to the fluctuation in salt concentration. Alternatively, the distinct nature of the environmental change in our study, which changes gradually in the case of the switch or transiently in the case of the pulse, compared with the instantaneous change of *Salingon et al*’s experimental design may be an important factor. This is consistent with prior work suggesting that gradually changing environments result in greater clonal interference than instantaneously changing environments in which mutations of large beneficial effect are more likely to fix early (Morley and Turner 2017).

### Genotypes exhibit distinct frequency dynamics

Whereas it has been widely demonstrated that cells exhibit rapid transcriptional (Gasch et al. 2000; Ronen and Botstein 2006; Airoldi et al. 2016; Spies et al. 2019) and physiological responses to changes in the environment (Bresson et al. 2020), the sensitivity of growth rate to environmental changes is less well studied. We sought to quantitatively describe the high resolution changes in genotype frequency across time for each genotype in each condition. The temporal dynamics of a genotype in a population is a result of both the externally imposed environmental selective pressure and the response to selection by all genotypes in the population. To characterize the dynamics of each genotype we performed hierarchical model fitting for each genotype using a model in which the log of the normalized barcode count from all ten time points is a polynomial function of time (**supplemental methods**). We explored the suitability of four distinct models of log normalized abundance versus time - linear, quadratic, cubic, and periodic. We quantified their applicability by starting with the most complex model and performing iterative model simplification using the log ratio of maximum likelihood test (**supplemental methods**).

We observed multiple distinct genotype dynamics. We find that between 10% - 30% of genotypes (**Figure 3F**) do not show a significant change in normalized abundance (**Figure 3A**) over the duration of the experiment. For these genotypes, the growth rates are not significantly different from the arithmetic mean over all genotypes. The prevalence of these genotypes is consistent with the greatest density of genotypes falling around a relative fitness of zero (**Figure 2C**). Although the mean relative growth rate is zero by our definition of relative growth rate, the concentration of the distribution around zero relative growth rate is not guaranteed or predictable. Many genotypes show log-linear behavior across time (**Figure 3B**) indicating sustained positive or negative selection. Whereas static conditions in which selection is constant may be expected to result in such behavior, we find that almost a quarter of all genotypes also exhibit log linear behavior in the switch condition (**Figure 3F**). Such genotypes that are unaffected by alternations in the environment may be indicative of generalists. We identified non-monotonic genotype dynamics in all three conditions (**Figure 3F**). Quadratic behavior (**Figure 3C**) indicates an accelerating or decelerating growth rate per cell, whereas cubic (or sigmoidal) behavior (**Figure 3D**) reflects two reversals in the sign of fitness over the course of the experiment. A similar diversity of behaviors is found in the two pulse conditions (**supplemental figure 9A**)

**Figure 3.**
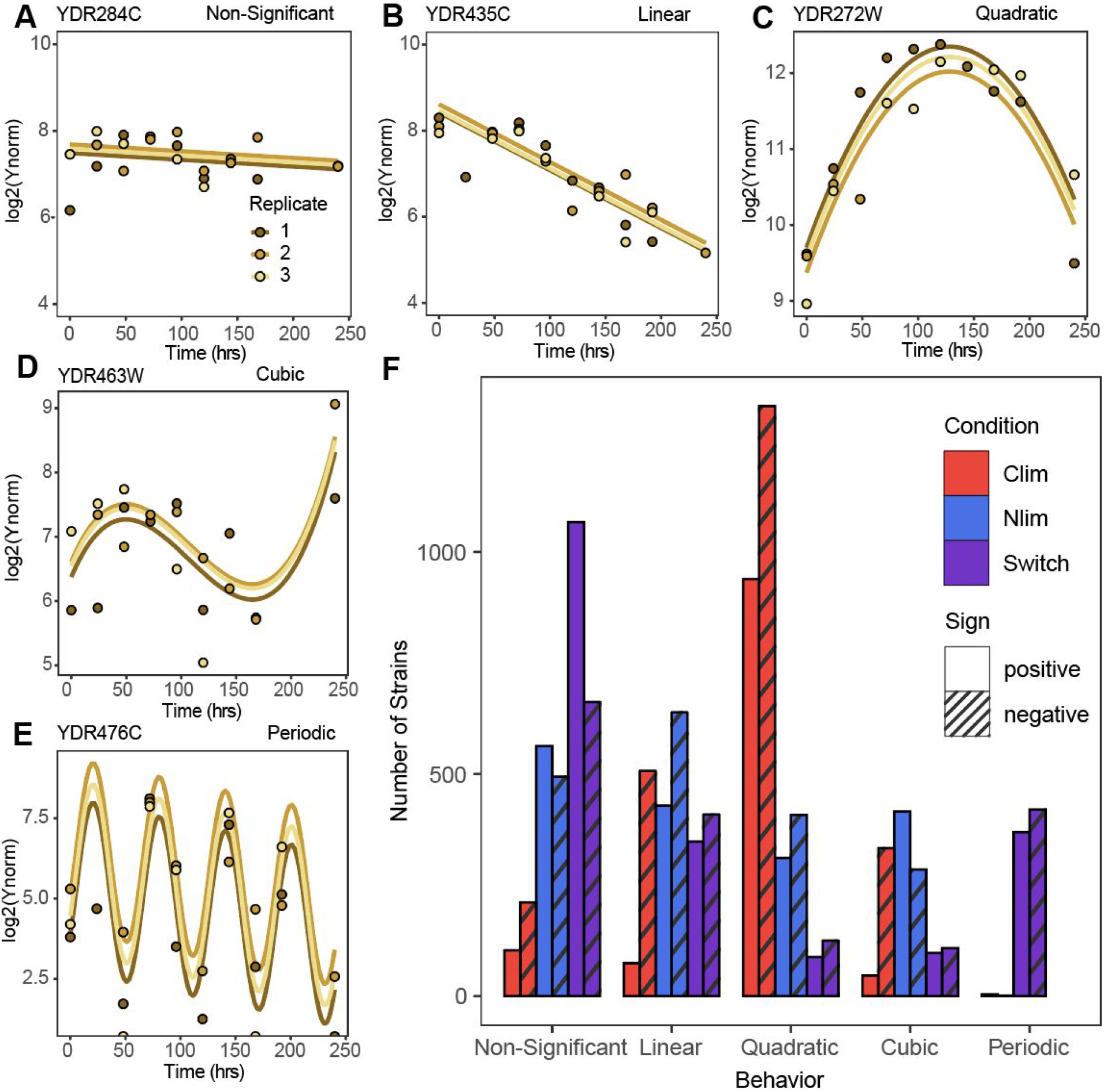
Diverse temporal trajectories of genotypes in different selective conditions. Temporal dynamics of genotypes across time fit to (**A**) non-significant, (**B**) linear, (**C**) quadratic, (**D**) cubic and (**E**) periodic models across time. (**F**) The distribution of temporal dynamics in each condition. Abundance changes are categorized as positive or negative based on the change in average growth rate between t = 0 and t = 240. Model fits for periodic models were defined as positive or negative on the basis of phase.

Our frequent sampling regime enables detection of genotype growth rate dynamics with high resolution. To that end, we tested whether genotypes show oscillatory behavior across the experiment (**supplemental methods**). We detect evidence for strong enrichment of periodically oscillating changes in genotype growth rate per cell (**Figure 3E**) that is unique to the switch condition (**Figure 3F** and **supplemental figure 9A**). In these genotypes the growth rate per cell oscillates with a period that matches the period of environmental change imposed by switching the feed media to the chemostat. These genotypes include both positive and negative phases (i.e. with 180 degree difference). This behavior suggests a class of genotype that is acutely sensitive to variation in the environmental condition. To the best of our knowledge, there are few cases in which such oscillations in genotype frequencies have been observed. One notable case is the oscillatory behavior of genotypes that has been observed over seasonal fluctuations in Drosophila populations ((Bergland et al. 2014; New et al. 2014), Machado et al. 2018). In addition, high resolution sequencing of the ‘Lenski lines’ identified genotype oscillations in evolving *E. coli* populations; however, this behavior eluded explanation (Good et al. 2017). Our finding suggests that such oscillations are potentially diagnostic of periodic variation in the environment. The 700 genotypes that comprise this class do not show significant enrichment for specific functions.

Non-monotonic behavior of genotypes may be the result of biological phenomena (e.g. environmental variation, genotype interactions, and density-dependent selection) or a consequence of data analysis methods. To test whether the highest frequency genotypes impact the apparent dynamics of other genotypes in the population, we computationally removed their barcodes from sequencing data, and repeated our complete analysis. We find that excluding the top performing genotype has a minimal effect on the identified non-monotonic growth behavior of the remaining genotypes (**supplemental figure 9B**). As expected, the same manipulation has drastic effects on diversity metrics in static conditions, but only a minimal effect on the results observed for fluctuating conditions (**supplemental figure 3**).

Fluctuating environments are enriched for genotypes that do not show a significant change in growth rate in comparison to other conditions (**Figure 3F** and **supplemental figure 9A**). This observation along with the unique oscillating genotypes point to two ways in which greater diversity is maintained in fluctuating conditions: 1) a larger fraction of genotypes have neutral fitness effects and 2) large fitness effects over short time spans undergo reversals in the direction of selection before they have a chance to dominate the population or go extinct.

### Environmental fluctuations select for specific mutant classes

To identify the biological pathways and processes that contribute to increased fitness in each condition we performed gene set enrichment analysis (GSEA) (Subramanian et al. 2005) using the ranked fitness measurements for each condition. We find that constant carbon limitation selection results in the positive selection of gene deletion mutants with functions in carbon metabolism (**supplemental figure 10**). The highest fitness genotype is deletion of *MTH1*, which has previously been reported as a target of selection in experimental evolution in glucose limited chemostats (Kvitek and Sherlock 2011). In static nitrogen limited chemostats, we find enrichment for genotypes with functions in nitrogen metabolism (**supplemental figure 10**). The highest fitness genotype is deletion of *GAT1*, which we have previously identified as conferring a fitness advantage in ammonium-limited chemostats (Hong and Gresham 2014; Hong et al. 2018). Interestingly, in our previous studies we identified *GAT1* hypomorphs as beneficial, but *de novo* null mutations in *GAT1* were not identified.

We identified enrichment for distinct gene functions that are unique to the switch condition. Specifically, deletions in genes that encode components of the endoplasmic reticulum associated degradation (ERAD) pathway including *HRD1*, *HRD3*, *USA1*, and *DER1* exhibit uniquely high fitness in the switch condition (**supplemental figure 11**). The ERAD complex is responsible for degrading misfolded proteins in the endoplasmic reticulum (ER). Loss of ERAD function may be uniquely beneficial in fluctuating conditions as decreased rates of protein degradation may facilitate persistence of proteins across conditions thereby serving as a type of ‘memory’ response.

The periodic addition of excess nutrients in the pulse conditions results in the enrichment of unique classes of genotype function in addition to functions that are shared with the static conditions (**supplemental figure 10**). This suggests that these transient environmental perturbations serve to both reduce the strength of selection and select for a unique class of genotypes.

### Fitness relationships between conditions

The fitness of a given genotype varies as a function of selection. We asked whether genotype behaviour under static selective conditions is predictive of fitness in a fluctuating environment. We find that the correlation between relative fitness in the two static conditions is low (**Figure 4A)**. The correlation between relative fitness in the carbon limited and switch condition (**Figure 4B)** and between the nitrogen limited and switch condition are somewhat higher (**Figure 4C**). We tested the simple model that fitness in a fluctuating environment is the mean of fitness in the two corresponding static conditions. We found that the correlation between the relative fitness in the switch condition and the mean of relative fitness in nitrogen limited and carbon limited conditions was only slightly increased compared with the correlation between each static condition and the switch condition fitness estimates (**Figure 4D**).

**Figure 4.**
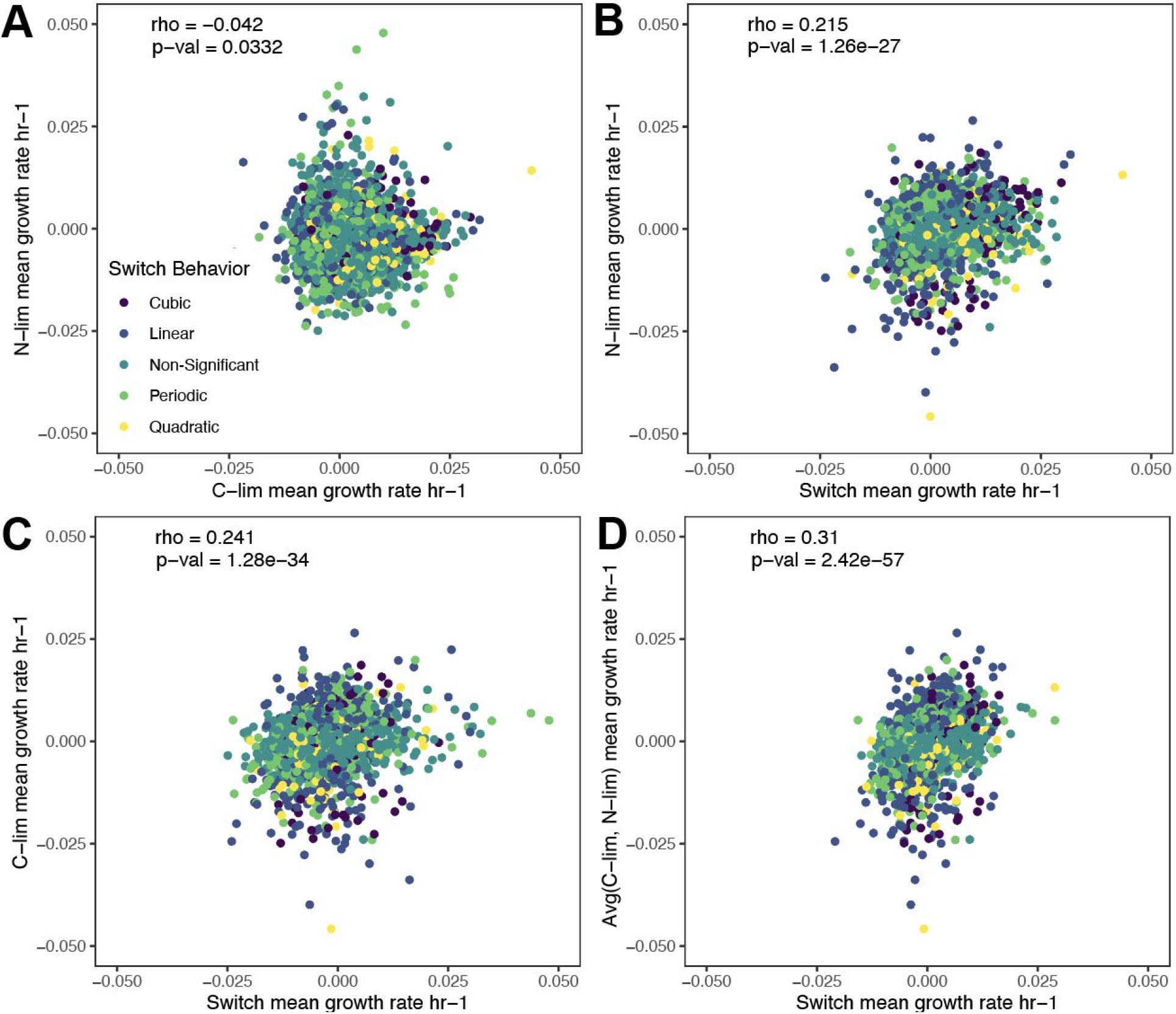
A subset of genotypes have a predictive relationship between fluctuating and static selective conditions. (**A**) The correlation in temporal mean growth rate per cell of genotypes between the two static conditions is low. There is intermediate correlation between the temporal mean growth rate per cell of the Switch condition and C-lim (**B**) and N-lim (**C**). The relationships between temporal mean growth rate per cell in the switch conditions and the average of the temporal mean growth rate per cell for the two static conditions has the highest correlation (**D**). Point colors indicate the model fit of the genotype as described in figure 3.

### Switching conditions harbor the highest short-term growth rates

To further understand how genotype behavior is affected in fluctuating conditions we compared short term fitness effects with long term fitness effects. Because we identified non-monotonic growth behavior, we calculated the piecewise fitness, defined as the mean relative fitness values between consecutive time points, in the static and switch conditions. We find that whereas the temporal average relative fitness across the full time course shows minimal differences in DFE between conditions (**Figure 2C)**, the piecewise DFE is highly dynamic between timepoints and conditions (**Figure 5A**). Whereas static conditions select for genotypes with the highest average growth rate across the full time course, the switching environment results in the largest short-term fitness values. We computed the variance in fitness at each time point and found that static conditions have a unique U-shaped variance pattern in contrast with the switch condition which showed oscillating piecewise fitness variance (**Figure 5B**). The large differences in variance in fluctuating conditions are explained by the behavior of the periodically oscillating genotypes which have the highest piecewise fitness values across all growth behaviors (**Figure 5C**). Periodically oscillating genotypes are not a uniform group as we identified four clusters of genotype behaviors. Three of the four clusters have unique overrepresented GO-terms suggesting functional coherence among these genotypes (**Figure 5D**).

**Figure 5.**
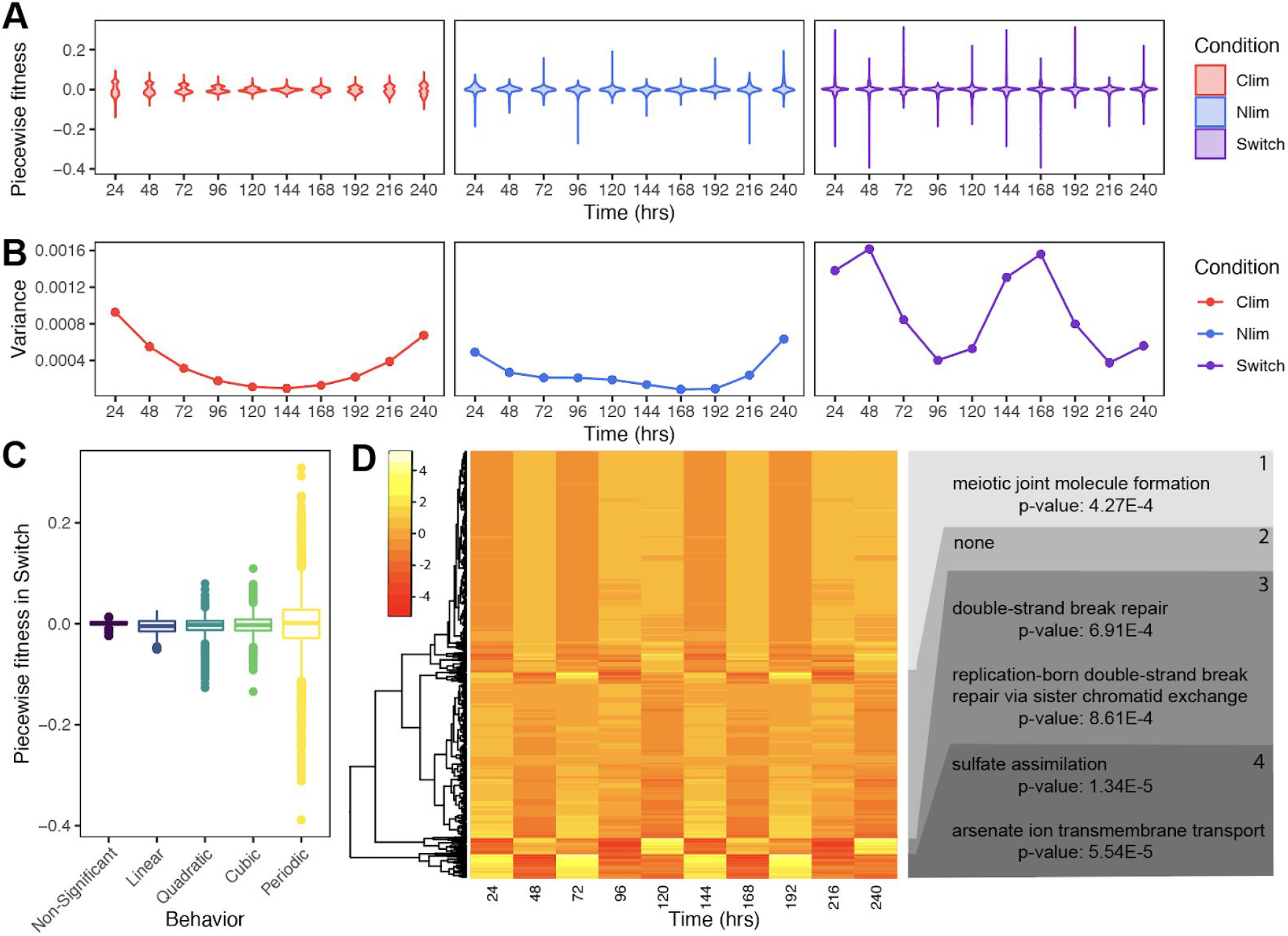
The switch condition uniquely results in short term fitness changes that are not detected over larger timescales. (**A**) Piecewise (temporal mean) relative fitness measurements were calculated by obtaining the difference between log normalized abundance at consecutive time points and dividing by the difference in time. Violin plots represent the distributions of piecewise fitness in each condition. (**B**) The variance of fitness measurements in each condition shows unique trends over time. (**C**) The distribution of piecewise fitness values according to best model fit. (**D**) Heatmap of scaled piecewise fitness for all periodically oscillating genotypes in the switch condition falling into four defined clusters. GO-terms that are enriched in each cluster are labeled on the right hand side.

## Discussion

In this study, we investigated the effect of fluctuating environments on genetic diversity and the distribution of fitness effects. We find that population diversity is greatly reduced in static environments compared with fluctuating environments. Our results support the idea that static environments impose stronger selection whereas fluctuating environments reduce the efficiency of selection (West and Mobilia 2020). By testing two distinct classes of environmental fluctuation we demonstrate that this result holds for two different types of environmental fluctuations.

We find that the maintenance of genetic diversity in fluctuating environments is a result of a combination of factors. First, genotypes with neutral fitness effects are enriched in fluctuating environments. Second, the presence of a unique class of genotypes that oscillate in frequency in fluctuating environments. Although this class includes genotypes with the highest and lowest short-term fitness effects the periodic reversal in the direction of selection ensures their maintenance at intermediate frequencies in the population, consistent with balancing selection. Third, the absence of genotypes with extreme long term fitness effects in fluctuating environments in contrast to static environments that are characterized by a single genotype with a large positive fitness effect that rapidly increases in frequency in the population. There has been considerable debate whether genetic diversity is primarily maintained through neutral fitness effects or through balancing selection (Hedrick et al. 1976). We have found that in the case in which new mutation does not occur, both balancing selection and neutral fitness effects contribute to the maintenance of genetic diversity in fluctuating environments.

Finally, we show that average fitness over long time spans can conceal the large variety of genotype behaviors in a population. Typically, fitness is estimated assuming monotonic behavior (Wiser and Lenski 2015) although a few studies have recently identified curvilinear dynamics (Schlecht et al. 2017). Our results suggest that the assumption of monotonic behavior is incorrect especially when considering population dynamics encompassing hundreds of unique genotypes, which is more representative of dynamics in natural populations (Wiser and Lenski 2015; Landis et al. 2021). This is the case even in static selective conditions. More complex selective regimes that result from environmental fluctuations can result in more complex genotype dynamics as illustrated by the unique class of oscillating genotypes identified in our study.

## Materials and methods

### Media

For all experiments, media consisted of defined minimal media supplemented with salts, metals, minerals, vitamins (Saldanha et al. 2004; Brauer et al. 2008; Airoldi et al. 2016). For glucose-limited media we added 0.08% glucose and 37mM ammonium-sulfate. For ammonium-sulfate-limited media we added 2% glucose and 400uM ammonium-sulfate. Static conditions used a single media source throughout the experiment. For the switch condition, we used a tube connecting the two feed media to a culture and alternated between the two media sources every 30 hours by manually clamping one inlet and opening the other. For pulse experiments we used the automated Sixfors chemostat system to deliver a bolus of either 40uM L-glutamine (PulseGln) or 40uM ammonium-sulfate (PulseAS) every three hours throughout the experiment.

### Experimental measurements of model parameters

Measurements were taken at time points 2.5 prior to switch, then at 17, 35, 44, 59.5, and 75.5 hours relative to the end of the first N-lim phase. This sampling scheme was chosen to capture the dynamics right after the first switch. Glucose was measured using the r-Biopharm Glucose kit. Ammonia was measured using the QuantiFluo™ Ammonia/Ammonium Assay Kit. Cell density and cell size was measured using a Coulter Counter.

### Mathematical modeling of chemostat growth in fluctuating environments

Population growth rate and the rate of change in the limiting nutrients glucose, and ammonium-sulfate were modeled using the following system of ordinary differential equations.

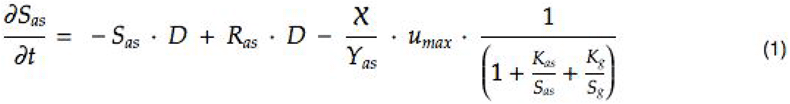

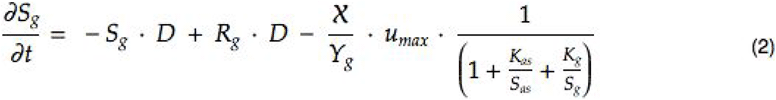

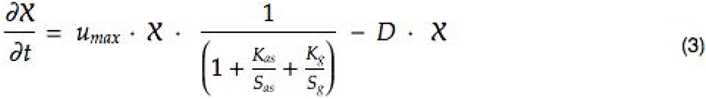

With the following parameters: D, the dilution rate of the culture (culture volumes/hr); X is the cell density (cells/mL), Y is the yield (cells/mL/mole of the limiting nutrient), u_max_ is the maximal growth rate constant (hr-1), R is the concentration (uM) of the limiting nutrient in the medium, and S (uM) is the growth limiting nutrient concentration in the chemostat. Equation (1) describes the changes in ammonium-sulfate concentration over time. Equation (2) describes the change in glucose concentration over time. Equation (3) describes the change in cell density over time. To study the effect of our experimental design for switching environments we performed numerical calculations with cell number (X) set to zero.

### Culturing conditions

Library construction was performed as described in (Sun et al. 2020). An aliquot (1.7×10^9^ cells/mL) of the pooled prototrophic gene deletion collection (VanderSluis et al. 2014) was thawed and 118uL were inoculated into triplicate chemostats with 200mL media for each condition. We estimate that this results in inoculation of the culture with 10^4 cells of each of the 10^3 genotypes. Cultures grew in r batch mode overnight at 30°C to allow cells to reach high density (3E7 cells/mL). The first sample was collected and then the media feed pumps were turned on and tuned to a dilution rate of 0.12^−hr^ to switch cultures to continuous growth. Samples were collected every 24 hours by passive sampling from the chemostat outlet for a total of 240 hours. Cell pellets were stored in −80C in a cell storage solution (0.9M sorbitol, 0.1M EDTA, 0.1M Tris). DNA extractions were performed using the Hoffman Winston DNA prep (Hoffman and Winston 1987). PCR amplification of barcodes of each sample was performed by using a universal primer and an indexed primer (Robinson et al. 2013). The P5 illumina adapter was incorporated to all samples. Barseq libraries were sequenced on a 1×75 bp run on a Illumina NextSeq500.

### Analysis of Barseq data

Barseq analysis was performed as previously described (Robinson et al. 2013). Briefly, barcode sequencing reads were matched to their corresponding genotypes using Barnone. Reads with base pair mismatches greater than 0 were excluded from the analysis. Libraries with less than 100,000 total read counts were removed (**Supplemental figure 1A**). Uptags and downtags for each genotype were summed and genotypes with aggregate counts across all conditions with less than 1000 were also removed (**Supplemental figure 1B**). DEseq2 was used to normalize libraries (Love et al. 2014). All code is available on github (https://github.com/far279/Abdul-Rahman_etal). Sequencing data has been submitted to the SRA.

### Mathematical modeling of genotype behavior

A detailed description of methods used for both data analysis and theoretical studies is provided in the supplemental methods. Throughout, we define the following terms:

- *Growth rate*: the change in population size between 2 time points, divided by time.
- *Instantaneous growth rate*: the derivative dn/dt.
- *Per capita (per cell) rate of change*: growth rate normalized by population size and accounted for by both cell divisions and cell death.
- *Per capita (per cell) growth rate*: same as per capita (per cell) rate of change where cell death is negligible.
- *Piecewise growth rate*: the growth rates between all consecutive timepoints based on the predicted values.

## Acknowledgements

We thank previous and present members of the Gresham, Vogel and Ghedin labs for valuable discussion and comments on the manuscript. We also thank Eugene Plavskin for comments on the manuscript and the NYU Genomics Core facility for sequencing services. This work was supported by NSF (MCB1818234) and NIH (R01GM107466, R01GM134066).

---

**Supplemental figure 1.**
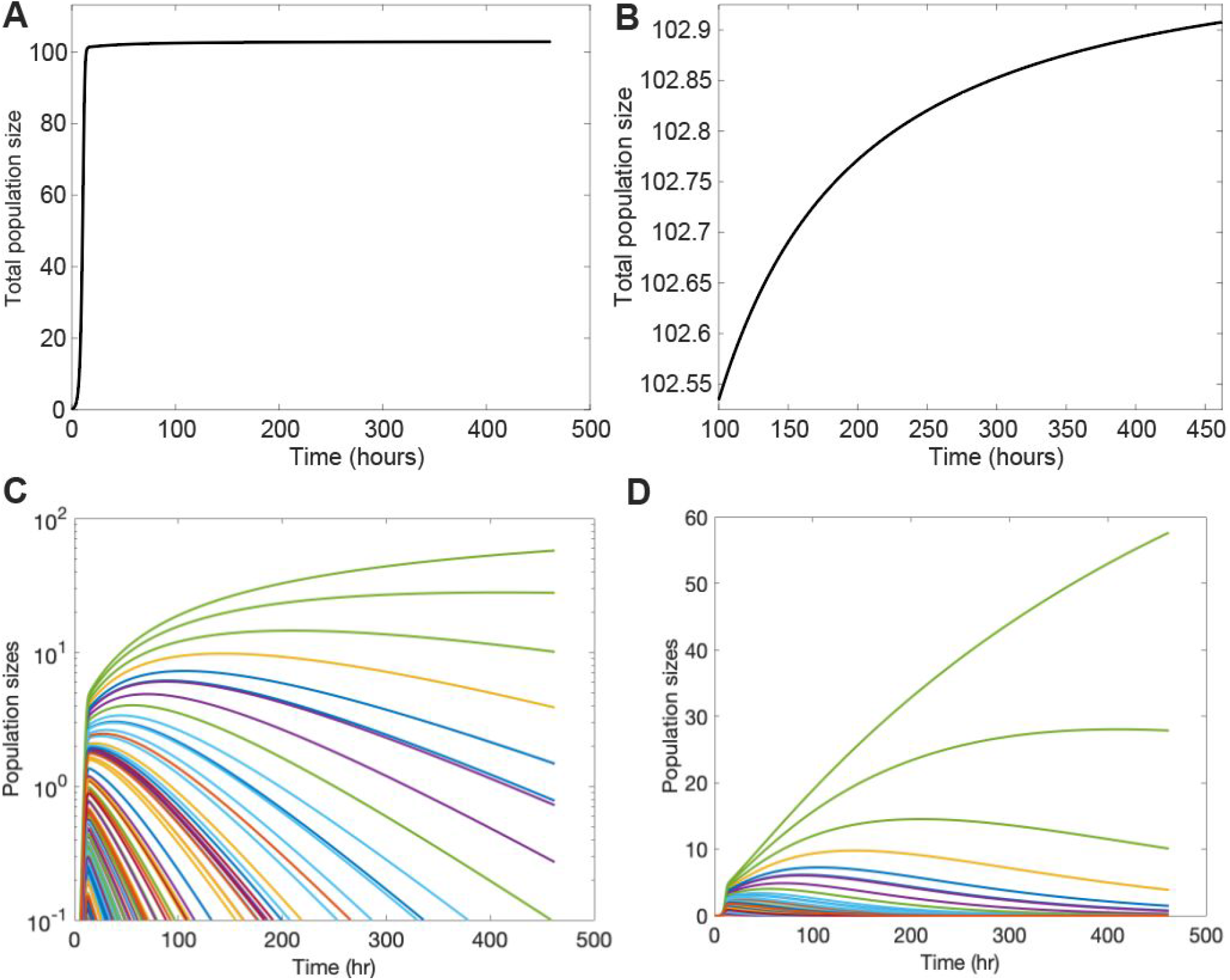
Modelling the growth of four thousand genotypes in the chemostat. **(A)** In the presence of thousands of genotypes the chemostat attains a quasi-steady state. **(B)** The total population size undergoes non-zero, but negligible, changes as selection acts on the population. **(C)** Individual genotype population sizes undergo large changes in frequency despite the relative invariance of total population size. (**D**) Dynamics of the top ten lineages.

**Supplemental figure 2.**
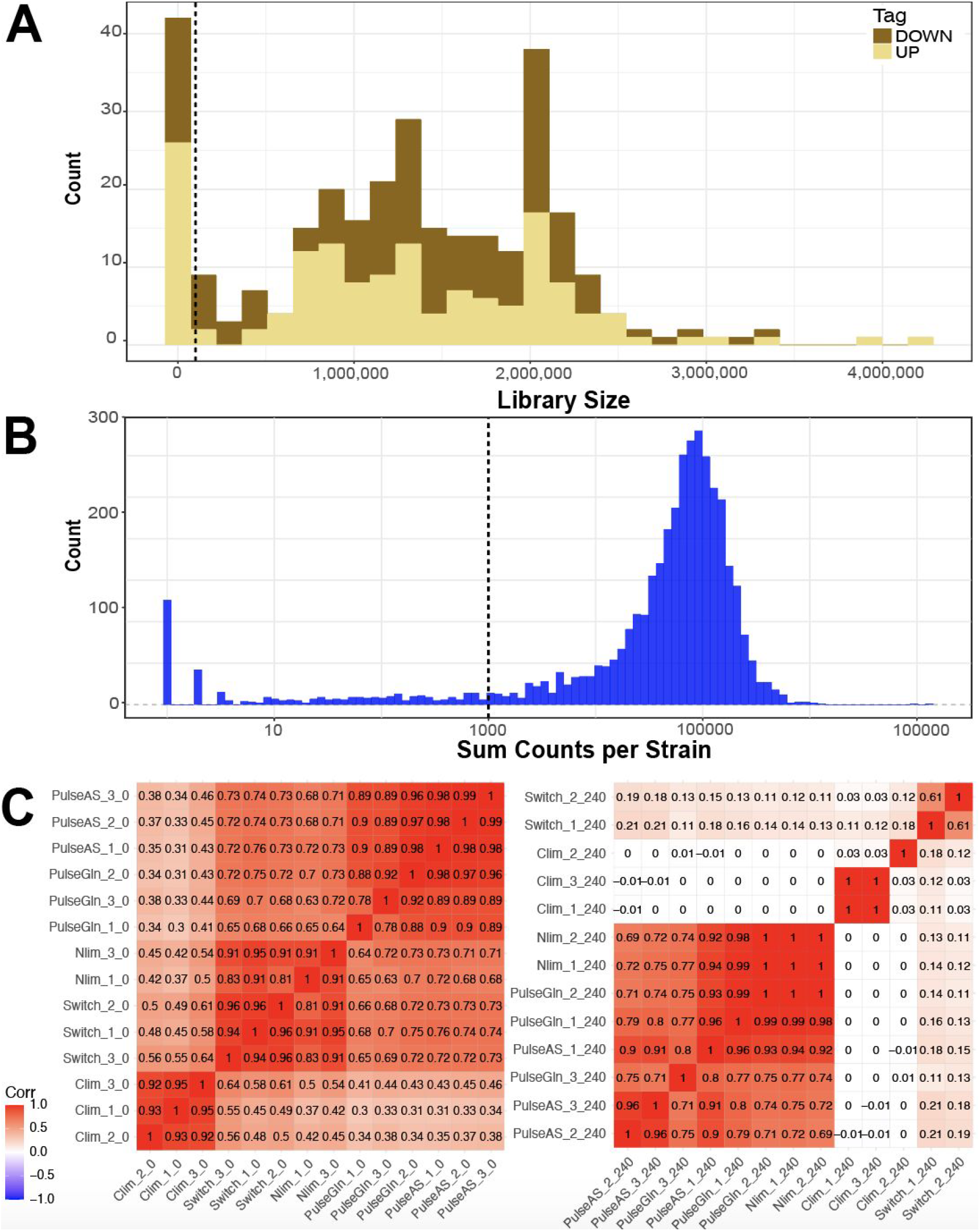
Barseq library quality control. (**A**) The complete distribution of library sizes before filtering is shown. The dashed line indicates libraries with less than 100,000 reads that were excluded from subsequent analysis. (**B**) The distribution of aggregate counts per genotype across all libraries (n =). The dashed line indicates genotypes with less than 1,000 aggregate reads that were excluded from subsequent analyses. (**C**) Pairwise Pearson correlation coefficients between all samples at the first (t = 0) (**left panel)** and final time point (t = 240) (**right panel**). Replicates with correlation coefficients less than 0.6 were removed before proceeding with analysis.

**Supplemental figure 3.**
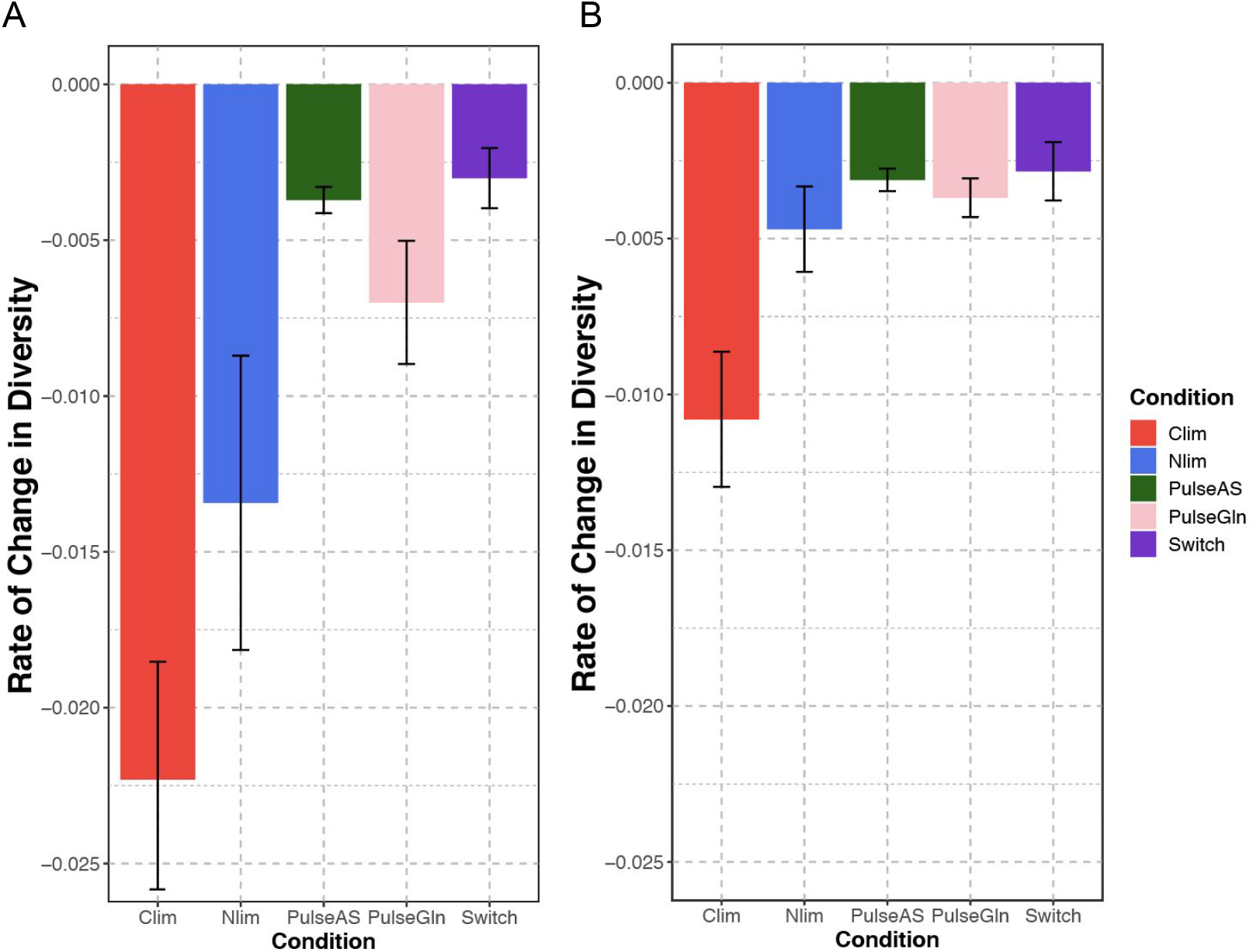
Maintenance of genetic diversity in different fluctuating selections. **(A)** Quantification of the dynamics of genetic diversity in other selective conditions. **(B)** Quantification of the dynamics of genetic diversity after excluding the highest fitness genotype in each condition.

**Supplemental figure 4.**
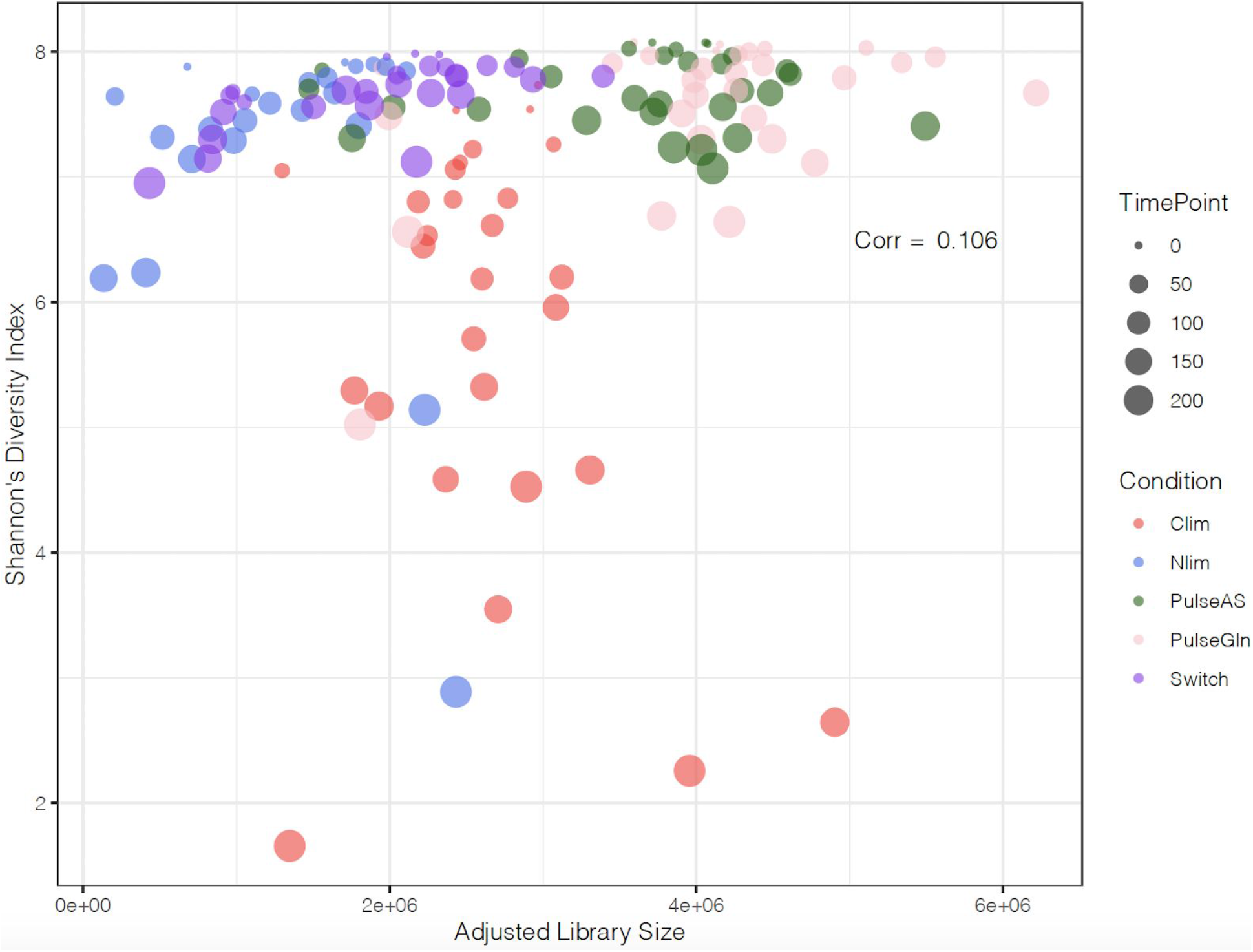
Library size does effect diversity estimates.

**Supplemental figure 5.**
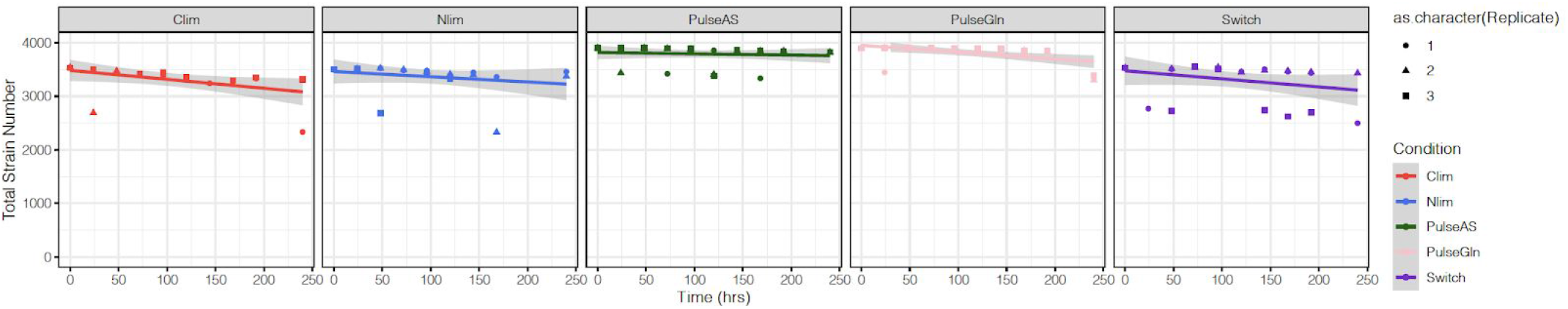
The rate of change in total strain number across all conditions.

**Supplemental figure 6.**
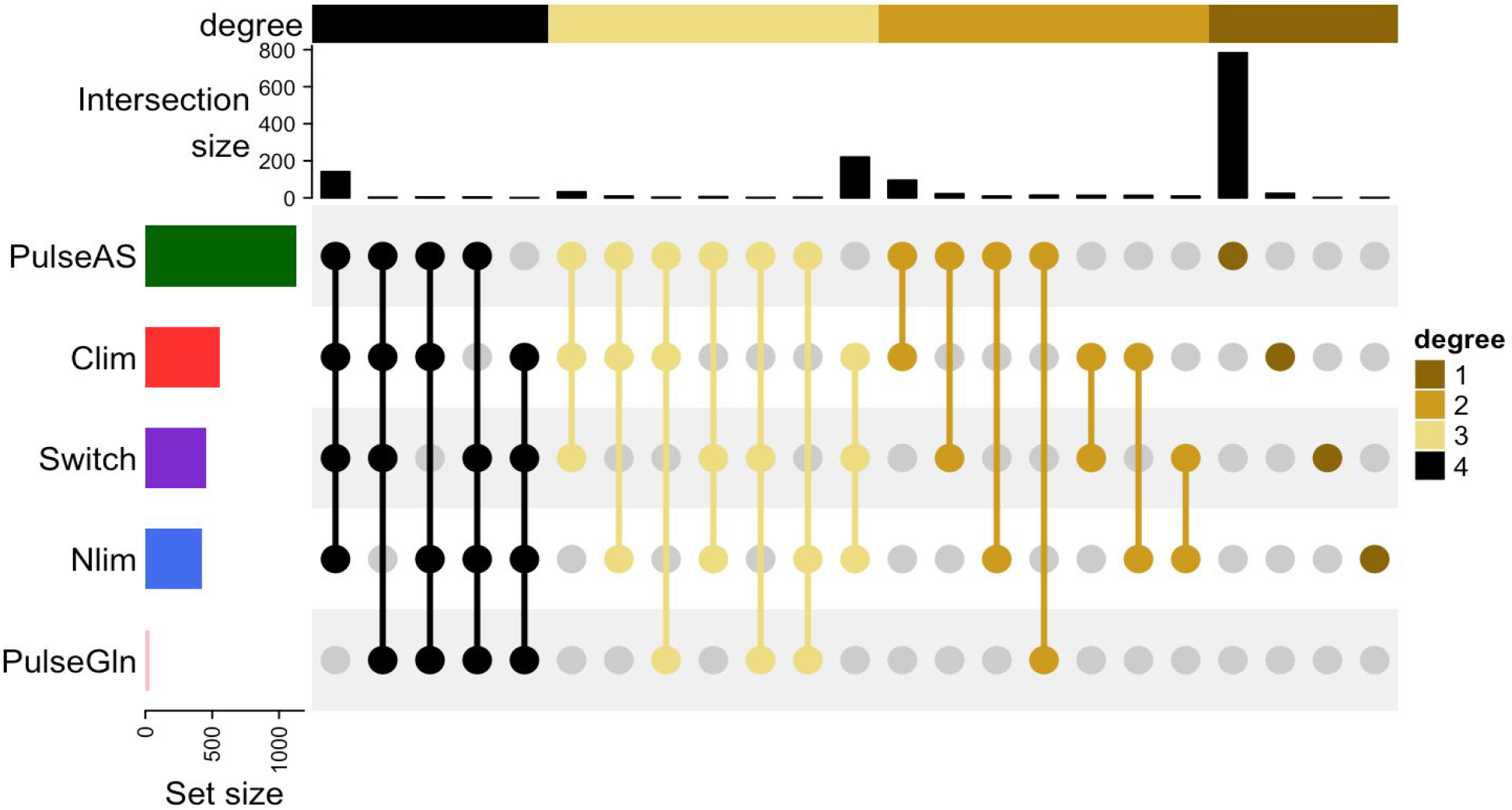
Strain extinction profiles in each condition. Extinct strains in the final time point (t = 240) are shared between subsets of conditions. Degree refers to the number of conditions that share a set of extinct genotypes.

**Supplemental figure 7.**
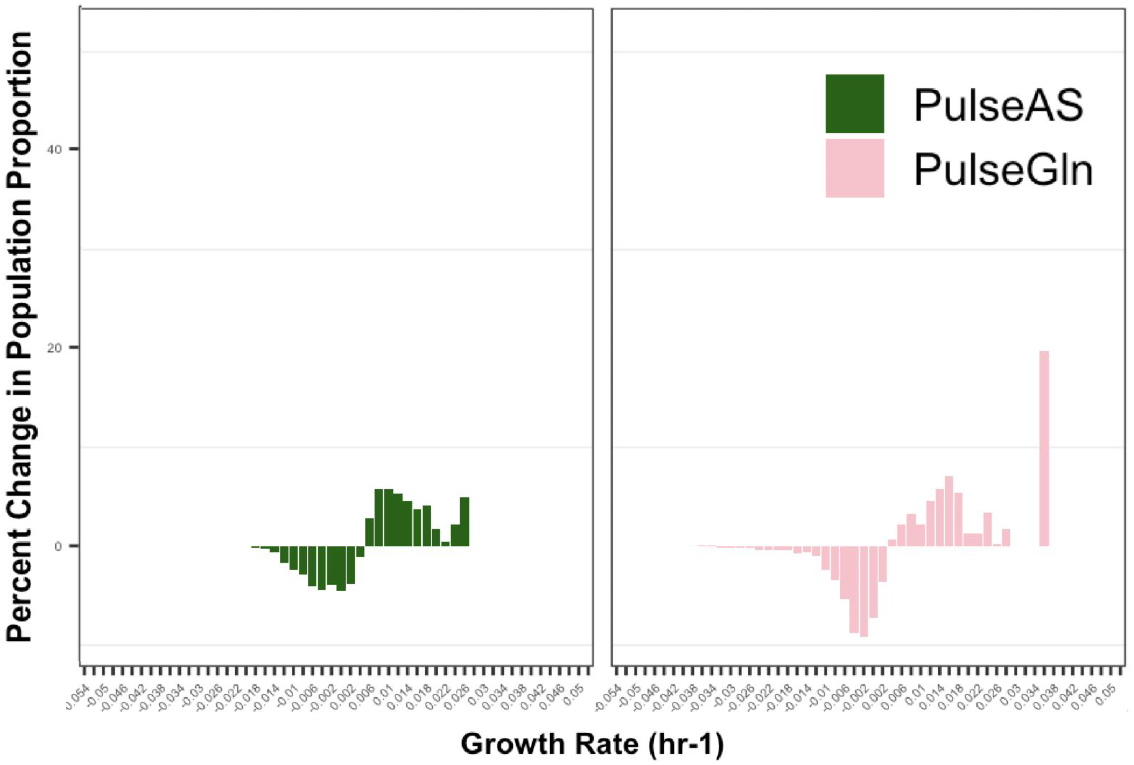
Change in percent population proportion for pulse conditions.

**Supplemental figure 8.**
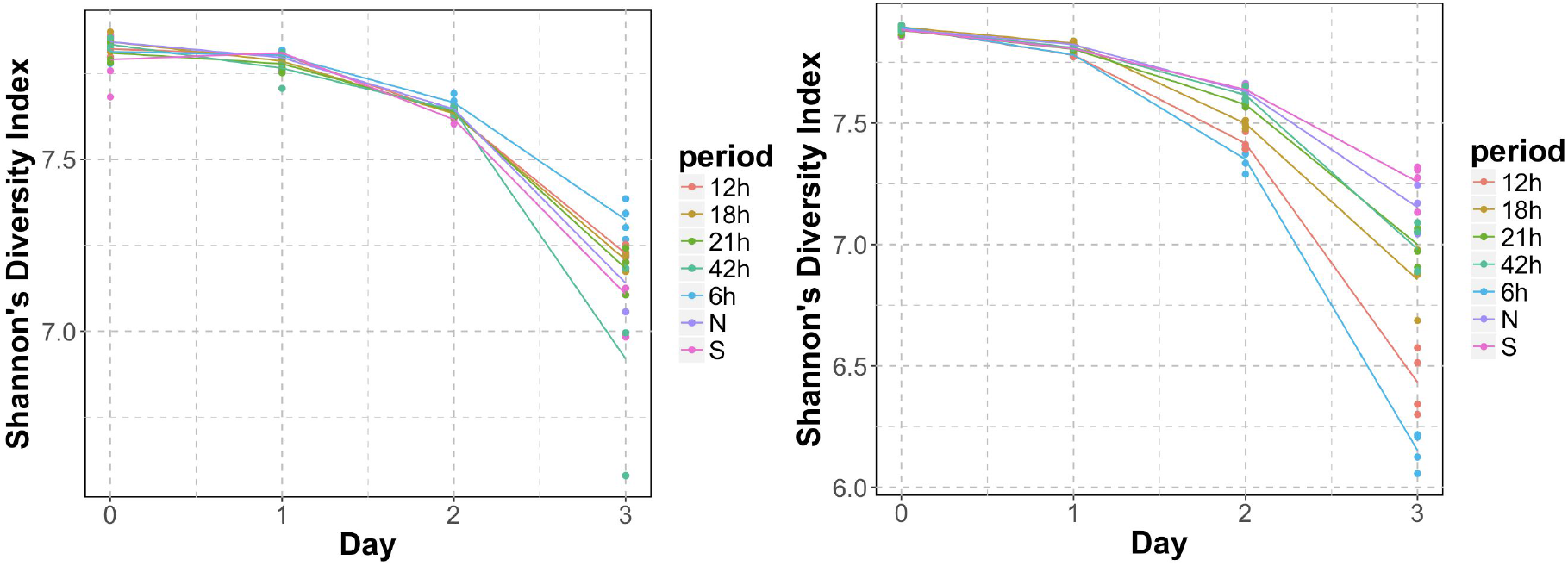
Diversity measurements of experiments from the Salingnon et al. dataset. Barseq was performed on the haploid gene deletion yeast library in conditions fluctuating between high (S) and low methionine (N) concentrations (left) and conditions fluctuating between salt (S) and no salt (N) concentrations (right).

**Supplemental figure 9.**
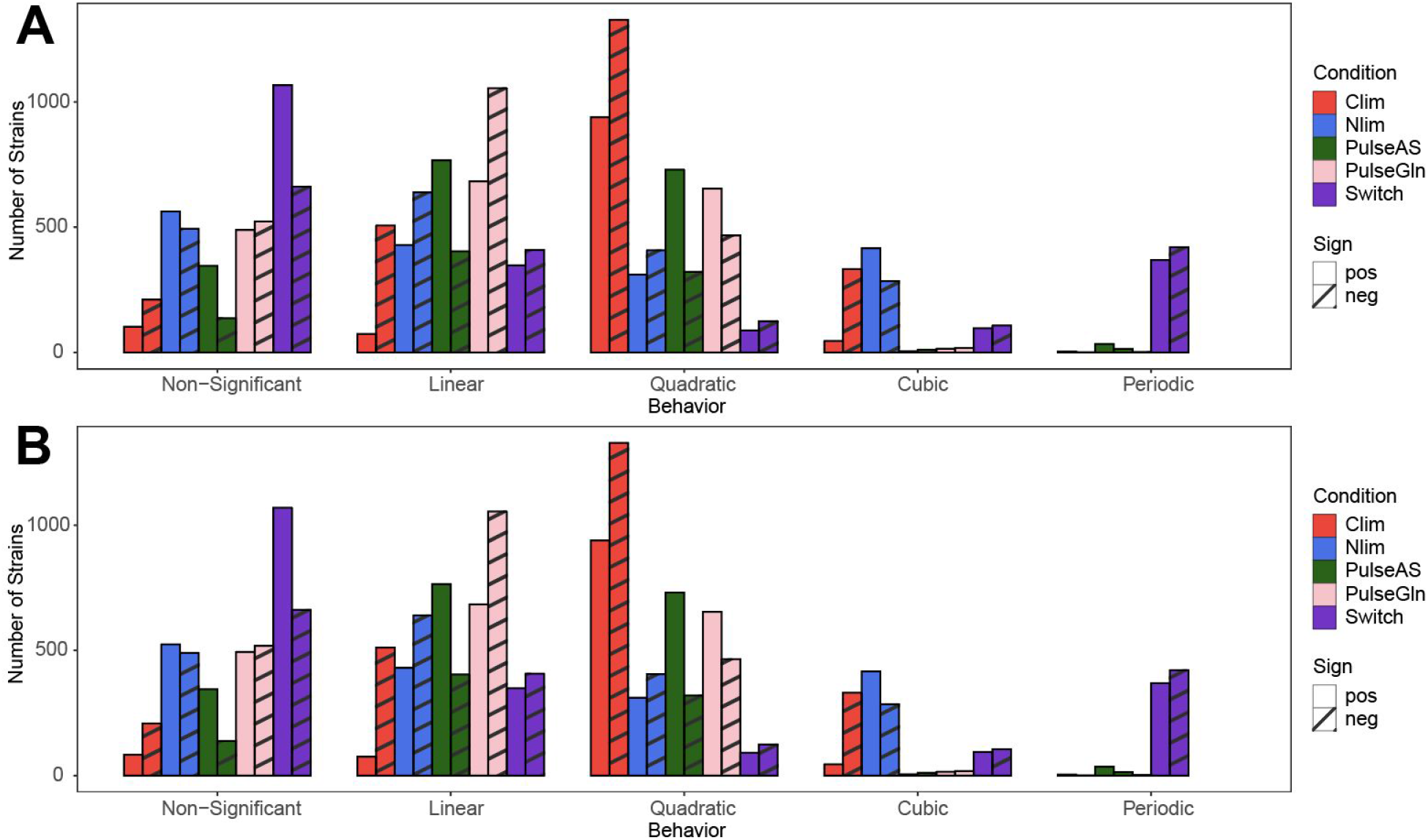
Genotype dynamics in static and fluctuating environments. **(A)** Extended summary of growth behavior including the two additional pulse conditions. **(B)** Reanalysis following removal of the highest fitness genotype does not alter the distribution of model fits.

**Supplemental figure 10.**
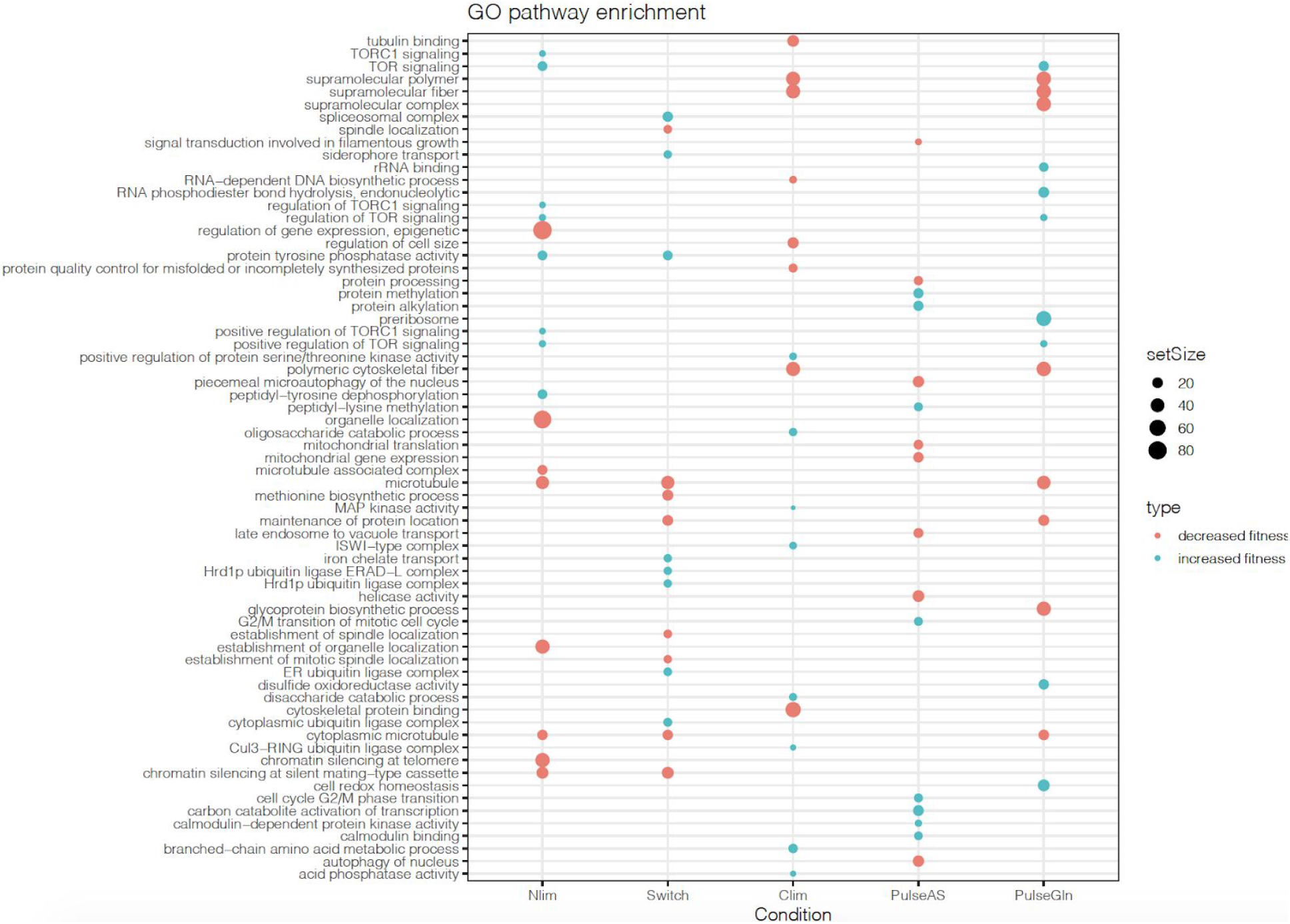
Gene set enrichment analysis (GSEA) of fitness effects in each condition. The top and bottom eight significantly (p-value < 0.05) represented GO terms for each condition are shown. Set size refers to the number of genes contained in a category.

**Supplemental figure 11.**
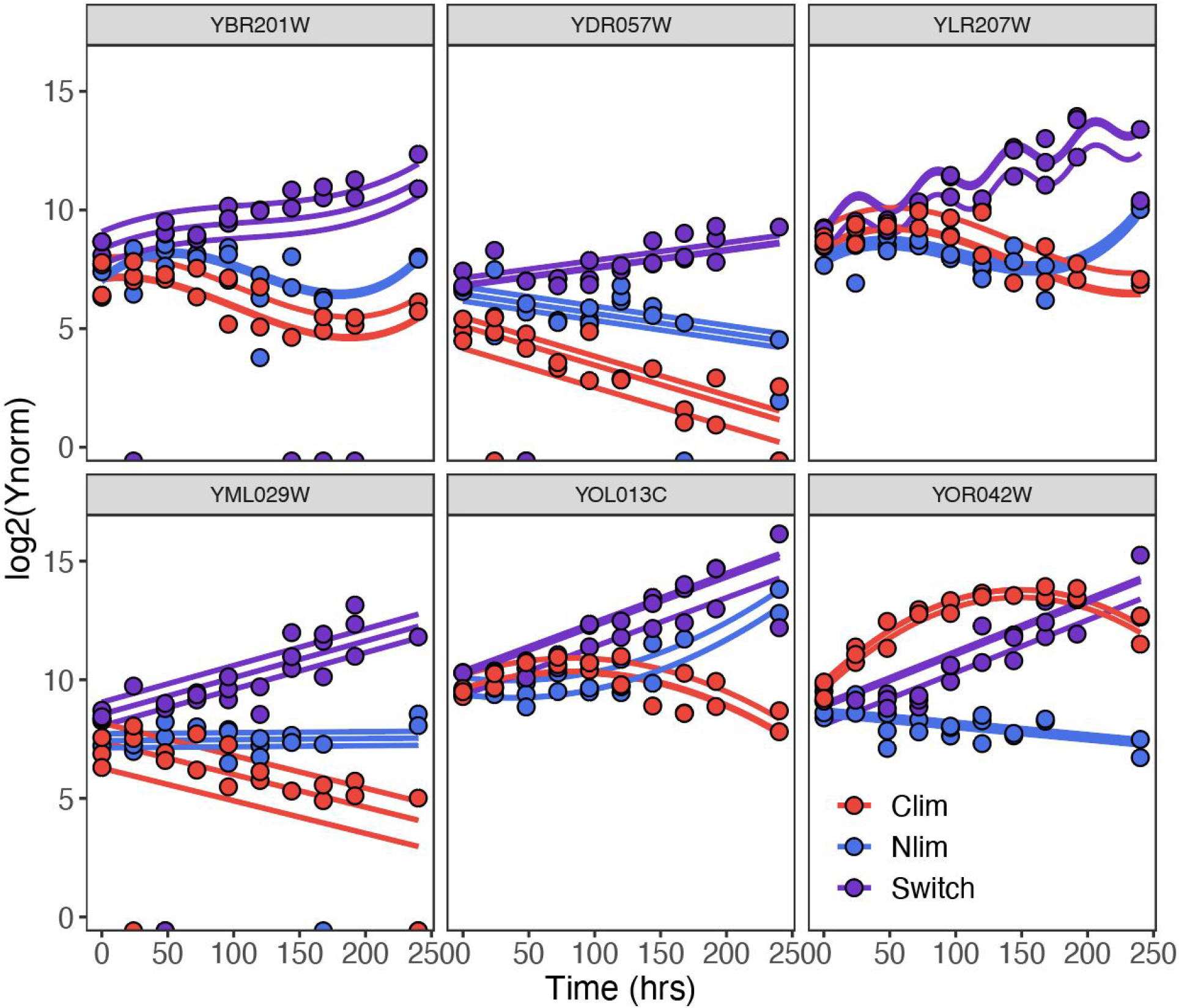
Deletion of the ERAD genes uniquely results in increased fitness in fluctuating environments. DER1, YOS9, HRD3, USA1, HRD1, and CUE5 gene deletions show consistent significant fitness increase in the switch condition but variable responses in carbon and nitrogen limiting conditions.t.

## SUPPLEMENTARY Tables

**Supplemental table 1. Pairwise correlation matrix of counts across all conditions.**

**Supplemental table 2.** DFE statistical measurements.

| Condition | Mode | Median | Mean | Max | Min | Range | Variance |
| --- | --- | --- | --- | --- | --- | --- | --- |
| Clim | 0.0077 | -0.0031 | -0.0027 | 0.0478 | -0.0271 | 0.0749 | 6.53E-05 |
| Nlim | 0.0124 | -0.0003 | -0.0007 | 0.0435 | -0.0439 | 0.0874 | 8.64E-05 |
| Switch | 0.0047 | 0.0005 | -0.0018 | 0.0265 | -0.0504 | 0.0769 | 9.53E-05 |
| PulseAS | 0.0020 | 0.0031 | 0.0027 | 0.0278 | -0.0176 | 0.0455 | 4.29E-05 |
| PulseGln | -0.0026 | -0.0004 | -0.0012 | 0.0361 | -0.0415 | 0.0776 | 6.94E-05 |

## Notes

### Competing Interest Statement

The authors have declared no competing interest.

