## Supplemental Methods for "Fluctuating environments maintain genetic diversity through neutral fitness effects and balancing selection"

February 2021

### S1 Mathematical and Statistical Methods

#### S1.1 Symbols and notation

Symbols, notation and mathematical expressions used in this section are summarized in Table S1.

#### S1.2 Basis of barcode count normalization

##### S1.2.1 Dynamics of genotypic populations

In the chemostat, a genetically homogeneous population of size $n$ grows through cell division at a rate given by $n'(t) = \lambda(t)\, n(t)$, where $\lambda(t)$ is the instantaneous, *time-dependent*, growth rate per cell; i.e., $\lambda(t) = n'(t)/n(t)$. We assume here that there is no cell death. If there were, $\lambda$ could be taken to mean the net rate of change per cell due to cell division and cell death. Each genotypic population in the chemostat is identified by its unique DNA barcode. Our normalization method allows us to estimate, for each genotypic population, the time average of its growth rate per cell (fitness), $\lambda_i(t)$, between any 2 time points, minus the arithmetic mean over all genotypes. Furthermore, with generalized linear model fitting, we will be able to estimate $\lambda_i(t)$ itself, minus the arithmetic mean. The normalization method is based on the population dynamics below.

In the chemostat, with $m$ genotypes, the number of cells with genotype $i$ at time $t$, $n_i(t)$, changes according to

$$\frac{d}{dt} n_i = [\lambda_i(t) - \beta]\, n_i, \text{ or} \tag{S1a}$$

$$\frac{d}{dt} \log(n_i) = \lambda_i(t) - \beta, \tag{S1b}$$

| Expression | Definition |
|---|---|
| $\lambda_i(t)$ | instantaneous growth rate per cell and our measure of fitness at time $t$ for sub-population with genotype $i$, as identified by its unique DNA barcode |
| $\Lambda_i(t) \,=\, \int_0^t \lambda_i(t')\, dt'$ | integrated fitness, for genotype $i$, over time interval $(0, t)$ |
| $(1/t) \int_0^t \lambda_i(t')\, dt'$ | temporal mean of fitness, for genotype $i$, over time interval $(0, t)$ |
| $m$ | number of genotypes in the chemostat |
| $\mu^{\mathrm{a}}_{\lambda}(t) \,=\, \frac{1}{m} \sum_{k=1}^{m} \lambda_k(t)$ | arithmetic mean, over genotypes, of instantaneous fitness at time $t$ |
| $\mu^{\mathrm{a}}_{\Lambda}(t) \,=\, \frac{1}{m} \sum_{k=1}^{m} \Lambda_k(t)$ | arithmetic mean of integrated fitness over time interval $(0, t)$ |
| $n_i(t)$ | size (or density) of sub-population with genotype $i$ at time $t$ |
| $N(t) \,=\, \sum_{i=1}^{m} n_i(t)$ | total number of cells (or density) in the chemostat at time $t$ |
| $p_i(t)$ | proportion of cells of genotype $i$ at time $t$ |
| $\lambda^{\mathrm{rel}}_i(t) \,=\, \lambda_i(t) \,-\, \mu^{\mathrm{a}}_{\lambda}(t)$ | relative (to arithmetic mean) fitness of genotype $i$ at time $t$ |
| $\Lambda^{\mathrm{rel}}_i(t) \,=\, \Lambda_i(t) \,-\, \mu^{\mathrm{a}}_{\Lambda}(t)$ | relative integrated fitness for genotype $i$ over time interval $(0, t)$ |
| $(1/t)\, \Lambda^{\mathrm{rel}}_i(t)$ | temporal mean of relative fitness for genotype $i$ over time interval $(0, t)$ |
| $\mu_{\lambda}(t) \,=\, \sum\limits_{i=1}^{m} \lambda_i(t)\, p_i(t)$ | population mean of fitness at time $t$ |
| $\sigma^2_{\lambda}(t) \,=\, \sum_{i=1}^{m} p_i(t)\, \left[\lambda_i(t) \,-\, \mu_{\lambda}(t)\right]^2$ | population variance of fitness at time $t$ |

Table S1: Definition of symbols and mathematical expressions

where $\beta$ is the dilution rate constant.

In our study, the growth rate per cell $\lambda_i(t)$ is empirical. Our experimental plots of log normalized barcode count versus time (Figure 3) show a rich variety of time courses in which many growth rates per cell are clearly not constant. We show below in our computational modeling (S1.6.2) that an extension of standard mathematical model for the chemostat [10] with heterogeneous populations [2], in which growth rate per cell depends instantaneously on the limiting nutrient concentration, $S(t)$, cannot account for the complex dynamics of cell numbers that we observe here (see Figure 3 and S1.6.2).

In our normalization method, the dilution rate constant $\beta$ plays no role. Ignoring $\beta$ and integrating Eq. (S1a) gives

$$n_i(t) \,=\, n_i(0)\, \exp\bigl\{\Lambda_i(t)\bigr\}, \text{ where} \tag{S2a}$$

$$\Lambda_i(t) \,=\, \int\limits_0^t \lambda_i(t')\, dt'. \tag{S2b}$$

Because $\Lambda_i(t)$ in Eq. (S2b) is the integral of the instantaneous fitness (growth rate per cell) for genotype $i$, $(1/t)\Lambda_i(t)$ is the temporal mean fitness over the time interval $(0, t)$.

The instantaneous fitness $\lambda_i(t)$ and the time average fitness $(1/t)\,\Lambda_i(t)$ are quantities we would like to estimate directly form our sequencing counts, but we don't know of any normalization method that can isolate these quantities. However, our normalization will allow us to determine for each genotype the fitness relative to the arithmetic mean over all genotypes. We define the relative fitness, $\lambda_i^{\mathrm{rel}}(t)$, as

$$\lambda_i^{\mathrm{rel}}(t) \stackrel{\mathrm{def}}{=} \lambda_i(t) - \frac{1}{m}\sum_{k=1}^{m}\lambda_k(t), \tag{S3}$$

and the time-average relative fitness, $(1/t)\Lambda_i^{\mathrm{rel}}(t)$, as

$$\frac{1}{t}\Lambda_i^{\mathrm{rel}}(t) \stackrel{\mathrm{def}}{=} \frac{1}{t}\left[\Lambda_i(t) - \frac{1}{m}\sum_{k=1}^{m}\Lambda_k(t)\right] \tag{S4}$$

##### S1.2.2 Dynamics of barcode tags

We assume that an effluent-sample from the chemostat that is used for sequencing has the same proportions of populations as those found in the chemostat.

We combine up- and down-tags [3] in our count of tags from each genotype. The proportion of barcode tags corresponding to genotype $i$ depends on all $n_i(t)$ and all "yield coefficients", $\alpha_k$, for $k = 1, 2, \ldots, m$, where $m$ is the total number of genotypes. The yield coefficient $0 \le \alpha_i \le 2$ can be thought of the average number of tags per cell contributed by a cell in population $i$. Based on our counts of up- and down-tags for each genotype in each sequencing run, and sometimes finding discrepant values, it appears that the $\alpha_i$ differ from DNA barcode to DNA barcode. In our method of normalization and inference relative genotype abundance across time, the $\alpha_i$ will play no role. However, they do contribute conceptually to the development of our normalization method.

The average average number of detected barcode tags per cell depends on the probability per cell of 2 non-disjoint events, $U_i$ — the up-tag from a cell in population $i$ is detected, $D_i$ — the down-tag is detected — and their complements, $U_i'$ and $D_i'$. One tag is captured in the event $(U_i \cap D_i')$; similarly, one tag is captured in the event $(D_i \cap U_i')$; and two tags are detected in the event $(U_i \cap D_i)$. Consequently, $\alpha_i = 1 \cdot P(U_i \cap D_i') + 1 \cdot P(D_i \cap U_i') + 2 \cdot P(U_i \cap D_i) = P(U_i) + P(D_i)$.

We assume that the number of sequencing tags from population $i$ in a sequencing sample depends on the proportion of detectable tags $i$ among the whole population of detectable tags in the sequenced sample, and on the library size $\mathcal{L}$.

With these assumptions, the expected number of barcode tags from population $i$ in a library at time $t$, $\mu_i(t)$, is given by

$$\mu_i(t) = \frac{\alpha_i n_i(t)}{\sum_{k=1}^{m}\alpha_k n_k(t)}\,\mathcal{L}. \tag{S5}$$

The library size, $\mathcal{L}$ whose estimate is not relevant in our normalization method. In analogy with RNA-seq methods [4, 5, 9], we model the random integer number of detected tags from population $i$, $Y_i$, as a negative binomial random variable with mean $\mu_i$ and a size parameter $a_i$, which is estimated in DESeq2 [4]. The motivation is that sampling noise alone would make the joint probability mass function of $[Y_1, Y_2, \ldots, Y_m]$ multinomial, which is very well approximated by a product of independent Poisson probability mass functions, as long as no particular genotype accounts for a sizable fraction of tags. However, in addition to the sampling noise, there are other sources of noise, such as the random number tags from populations of low number, stemming from the random number of such cells, deviations of $\alpha_i$ from the population average from replicate to replicate for fixed $i$, and experimental noise. In this case, the additional sources of noise imply that an overdispersed Poisson model is in order. The obvious choice, in analogy with RNA-seq, is the negative binomial.

#### S1.3 Normalization of barcode counts

We propose a normalization of counts, $y_i$, for genotype $i$ in a library, by a factor, $s(t)$, that is given by the geometric mean of counts over all genotypes in the library. This normalization method does not give a quantity for each genotype that is directly related to its proportion in the overall population. Such a normalization method is not appropriate for inference of instantaneous growth rates per cell (fitness), or time-average fitness. The reason is that, while the numerator term in a proportion is determined in simple way as a function of time that is dependent on the instantaneous fitness, the denominator term is a function of time that depends on each of the individual population sizes as a function of time, rather than constant.

As shown below, the utility of our normalization is that it allows one to easily compute, for each genotype $i$, the time average fitness between any 2 time points, minus the arithmetic mean of this quantity over all genotypes in Eq. (S4). We refer to this as the relative temporal-mean fitness, where the term *relative* signifies relative to the arithmetic mean over genotypes. The same sort of definition of terms applies to instantaneous fitness and instantaneous relative fitness.

The reasoning behind our normalization stems from the fact that the geometric mean, over all genotypes, of population sizes, changes as an exponential function of time. To illustrate this idea, we consider the special case of growth rate constants per cell that are independent of time, and we ignore dilution. The well known argument is as follows

$$\left[\prod_{j=1}^{m}\frac{n_j(t)}{n_j(0)}\right]^{1/m} = \left[\prod_{k=1}^{m}\exp\{\lambda_k\, t\}\right]^{1/m} \tag{S6a}$$

$$= \left[\exp\left(\sum_{k=1}^{m}\lambda_k\, t\right)\right]^{1/m} \tag{S6b}$$

$$= \exp\left(\left[\frac{1}{m}\sum_{k=1}^{m}\lambda_k\right]\, t\right) \tag{S6c}$$

$$= \exp\left(\mu_\lambda^{\mathrm{a}}\, t\right), \tag{S6d}$$

where $\mu_\lambda^{\mathrm{a}}$ is defined implicitly by Eqs. (S6c) and (S6d) as the arithmetic mean of $\lambda_k$ over all $k$ (genotypes).

In the case where we have time-dependent growth rates per cell, $\lambda_i(t)$, and dilution rate, $\beta$, the geometric mean of the population sizes is given by

$$\left[\prod_{k=1}^{m}\frac{n_k(t)}{n_k(0)}\right]^{1/m} = \exp\left(\frac{1}{m}\sum_{k=1}^{m}\Lambda_k(t)\, -\, \beta\, t\right) \tag{S7a}$$

$$= \exp\left(\mu_\Lambda^{\mathrm{a}}(t)\, -\, \beta\, t\right), \tag{S7b}$$

where $\mu_\Lambda^{\mathrm{a}}(t)$ is defined implicitly by Eqs. (S7a) and (S7b) at the arithmetic mean of $\Lambda_k(t)$ over all $k$. Recall that $\Lambda_k(t)$ is the integrated fitness of population $k$ over the time interval $(0, t)$.

What do we get when we normalize counts $y_i(t)$ by

$$s(t) \stackrel{\mathrm{def}}{=} \left[\prod_{k=1}^{m} y_k(t)\right]^{1/m}, \tag{S8}$$

the geometric mean over $k$ of counts $y_k(t)$? To answer this question, first, we treat the normalization factor, $s(t)$, as a known parameter, rather than a random variable. This is the customary treatment of normalization factors in high throughput sequencing (e.g., DESeq2 [4], edgeR [5, 9], and limma [8]). Second, for sake of exposition only, we go further and replace $s(t)$ by the geometric mean over $k$ of the expected values of the counts, $\mu_k(t)$, rather than the actual counts. With this normalization, Eq. (S2a) for $n_i(t)$, and Eq. (S5) for

$\mu_i(t)$, the normalized count simplifies, and it has an expected value given by

$$\frac{\mu_i(t)}{s(t)} \approx \frac{\mu_i(t)}{\left[\prod_{k=1}^{m} \mu_k(t)\right]^{1/m}} \tag{S9a}$$

$$= \frac{\alpha_i \, n_i(t)}{\left[\prod_{k=1}^{m} \alpha_k \, n_k(t)\right]^{1/m}} \tag{S9b}$$

$$= \frac{\alpha_i \, n_i(0)}{\left[\prod_{k=1}^{m} \alpha_k \, n_k(0)\right]^{1/m}} \exp\left[\Lambda_i(t) \,-\, \mu_\Lambda^{\mathrm{a}}(t)\right] \tag{S9c}$$

$$= \exp\left(\gamma_i + \Lambda_i(t) \,-\, \mu_\Lambda^{\mathrm{a}}(t)\right), \tag{S9d}$$

where

$$\gamma_i \stackrel{\mathrm{def}}{=} \log \frac{\alpha_i \, n_i(0)}{\left[\prod_{k=1}^{m} \alpha_k \, n_k(0)\right]^{1/m}} \tag{S10}$$

is the log of the normalized count at $t = 0$.

###### S1.3.1 Time-average relative fitness and inter-population differences between time-average fitness

It is convenient to give the expected normalized count for population $i$ a name,

$$z_i(t) \stackrel{\mathrm{def}}{=} \frac{\mu_i(t)}{s(t)}. \tag{S11}$$

From Eq. (S9d) one can see that the time-average relative fitness over time interval $(0, t)$, as defined in Eq. (S4), is given by

$$\begin{aligned} \frac{1}{t}\Lambda_i^{\mathrm{rel}}(t) &\stackrel{\mathrm{def}}{=} \frac{1}{t}\left[\Lambda_i(t) \,-\, \mu_\Lambda^{\mathrm{a}}(t)\right] \\ &= \frac{\log z_i(t) \,-\, \log z_i(0)}{t}. \end{aligned} \tag{S12}$$

Similarly, the time-average relative fitness over any time interval $t_1 < t < t_2$ is given by

$$\frac{\log z_i(t_2) \,-\, \log z_i(t_1)}{t_2 \,-\, t_1},$$

the difference of log normalized count at times $t_2$ and $t_1$, divided by the time difference. The instantaneous relative fitness for genotype $i$ is given by

$$\lambda_i^{\mathrm{rel}}(t) \,=\, \frac{d}{dt} \log z_i(t). \tag{S13}$$

One can compute the difference between the time-average fitness for 2 genotypes, $i$ and $j$, from their relative time-average fitness (because the arithmetic average term is common to all genotypes in the chemostat):

$$\frac{1}{t}\left[\Lambda_i(t) \,-\, \Lambda_j(t)\right] \,=\, \frac{1}{t}\left[\Lambda_i^{\mathrm{rel}}(t) \,-\, \Lambda_j^{\mathrm{rel}}(t)\right] \tag{S14}$$

#### S1.4 Polynomial modeling

We found that many of our normalized bar counts could not be accounted for by a model in which the growth rate per cell is constant over time. This would not be surprising if the thousands of interacting genotypes in the chemostat at once condition the growth medium beyond affecting the concentration of the common limiting nutrient, $S(t)$. If so, we expect a variety of time courses of $\lambda_i(t)$. One can think of these behaviors, in the context of the standard chemostat model [10], as an extension in which the maximum growth rate per cell, $\lambda_i^{\max}$, and the half-saturation constant, $K_i$ in the Monod growth rate function, are both functions of times, so that

$$\lambda_i(t) \;=\; \lambda_i^{\max}(t)\,\frac{S(t)}{K_i(t)\,+\,S(t)}. \tag{S15}$$

Our informal inspection of many normalized bar counts as a function of time suggested the possibility that the log of normalized counts could be fit by simple polynomial functions in $t$. Consequently, we modeled the exponential argument in Eq. (S9d), for the normalized count,

$$\gamma_i \,+\, \Lambda_i(t) \,-\, \mu_\Lambda^{\mathrm{a}}(t),$$

by a polynomial in $t$, up to $3^{\mathrm{rd}}$ order; i.e,

$$\frac{\mu_i(t)}{s(t)} = \exp\left(\gamma_i \,+\, \Lambda_i(t) \,-\, \mu_\Lambda^{\mathrm{a}}(t)\right) \tag{S16}$$

$$\approx \exp\left(b_0 + b_1 t + b_2 t^2 + b_3 t^3\right) \tag{S17}$$

In the absence of terms higher than first order in the exponential argument in Eq. (S17), the coefficient $b_1$ is the fixed relative growth rate per cell in population $i$, $\lambda_i^{\mathrm{rel}}$.

We used DESeq2 to compute the $b$-coefficients in Eq (S17). Eq. (S17) is a standard generalized linear model with a log link function. Recall that we assume that the random barcode count $Y_i(t)$ is distributed as a negative binomial random variable with mean $\mu_i(t)$ and shape parameter $a_i$ (inverse of *dispersion*, as the term is used in DESeq2 [4]). We started our model fitting with the $3^{\mathrm{rd}}$ order model and performed sequential model simplification by comparing the order-$k$ model to the order-$(k-1)$ model. For the comparison, we used the log ratio of maximum likelihoods test in DESeq2 [4], which gives a $p$-value and an adjusted $p$-value (False-Discovery Rate, FDR) for each simplification. All simplifications with an adjusted $p$-value greater than 0.1 were accepted in each round of simplifications. If $b_k$ was found to be significant, we did not test model simplifications at levels below $k$. The reason is that, if orthogonal polynomials [6] in $t$ were used as the basis for polynomial modeling, rather than powers of $t$, the orthogonal polynomial with highest order $k$, would, in general, include all powers less than $k$.

#### S1.5 Fourier modeling for periodic environments

For our experiments with fluctuating environments, the chemostat input was switched periodically, in a square-wave manner, between two limiting nutrients, with period $T = 60\,\text{hr}$. The corresponding frequency of the switching is $f = 1/T$. If we assume that the two limiting nutrients in the chemostat also fluctuate periodically at with same period, as confirmed in Fig. 1C, we expect to find some genotypes whose instantaneous growth rates per cell respond with periodic fluctuations, also with the same period. According to Fourier theory, a periodic function of $t$ for $0 < t < T$ can be expressed as a sum of sinusoidal components at integer multiples of the fundamental (input) frequency, $f = 1/T$.

We explored a model in which we assumed that the dynamics underlying $\lambda_i(t)$ are slow enough, so that $\lambda_i(t)$ is well approximated by a constant plus a single sinusoidal component at the fundamental frequency, $f = 1/T$. A mathematical formulation for this class of models is

$$\frac{1}{n}\frac{d}{dt}n = \lambda_0 + \lambda_1 \cos(2\pi f t + \theta), \qquad \text{or} \tag{S18a}$$

$$\frac{d}{dt}\log n = \lambda_0 + \lambda_1 \cos(2\pi f t + \theta), \tag{S18b}$$

where $\lambda_0$ and $\lambda_1$ are coefficients to be fit, and $\theta$ is the phase angle of the sinusoid. The solution to Eq. (S18b) is

$$\begin{aligned} \log\frac{n(t)}{n(0)} &= \lambda_0\, t \,+\, \lambda_1 \frac{1}{2\pi f}[\sin(2\pi f t + \theta) \,-\, \sin\theta\,] \\ &= -\frac{\lambda_1}{2\pi f}\sin\theta + \lambda_0\, t \,+\, \lambda_1 \frac{1}{2\pi f}\,\sin(2\pi f t + \theta) \end{aligned} \tag{S19}$$

Consequently we modeled the argument of the exponential in the equation for $\mu_i(t)/s(t)$ (expected value of normalized count) Eq. (S9d), for fluctuating environments, as

$$\gamma_i \,+\, \Lambda_i^{\text{rel}}(t) \;\approx\; b_0 + b_1 t + b_2 \sin(2\pi f t) + b_3 \cos(2\pi f t). \tag{S20}$$

In Eq. (S20) we have used the trigonometric identity,

$$b_2 \sin(2\pi f t) + b_3 \cos(2\pi f t) \;=\; \sqrt{b_2^2 + b_3^2}\,\sin(2\pi f t + \theta),$$

where $\sqrt{b_2^2 + b_3^2}$ is the amplitude of the sinusoidal component of the response, and $\theta$ is the phase lag (angle) relative to the fundamental component of the periodic switching of the input limiting nutrient. The phase lag $\theta$ is given by $\arctan(b_2/b_3)$. We computed $\arctan(b_2/b_3)$ from the fitted parameters $b_2$ and $b_3$ by using the *atan2* function in R, which keeps track of the quadrant in which $\theta$ lies.

We tested the null hypothesis of no sinusoidal component (log maximum likelihood test), and we used the same FDR as that used in the polynomial modeling. We found roughly 700 out of roughly 4000 genotypes with significant sinusoidal fluctuation in the instantaneous growth rate per cell (minus the arithmetic mean). See Fig. 3E for an example of such a genotype.

Failure to reject the null hypothesis for a genotype made that genotype a candidate for a polynomial model. For sake of simplicity, we did not consider mixtures of $2^{\text{nd}}$ or $3^{\text{nd}}$ degree polynomials with an additional sinusoidal component.

#### S1.6 Expectations for a chemostat with heterogeneous populations and fixed limiting-nutrient input: theory and computational modeling

##### S1.6.1 Theoretical results on the possibility of fixed cell number and the coexistence of several genotypes with differing growth rate constants in the steady state

Our experimental results, with steady influx of a single limiting nutrient to the chemostat, show that, after a brief transient period before our experimental clock time $t = 0$, both the total cell number, $N(t)$, and the concentration of limiting nutrient, $S(t)$ (Figure 1D, left panel), appear to be constant. Meanwhile, the population proportions evolve, with decreasing Shannon diversity index (Figure 2B). At $t = 240\,\text{hr}$, a single genotype accounts for large proportion of total cells (Figure 2C). Because our experimental time period was only 240 hr (roughly 40 generations), one might inquire about the ultimate fate of cell number and population proportions, assuming that a steady state is achieved.

Although a straightforward extension of the basic chemostat model [10] for numerous heterogeneous populations (genotypes) in the chemostat [2] cannot reproduce the rich dynamical behavior we see in our experiments (S1.6.2), it is helpful to state the model equations here as a way of anchoring our thoughts. In the case of heterogeneous populations, the chemostat equations become,

$$\frac{d}{dt}N = \left[\mu_\lambda(t) - \beta\right] N \tag{S21a}$$

$$\frac{d}{dt}S = (S_0 - S)\,\beta - \frac{N}{Y}\mu_\lambda(t) \tag{S21b}$$

$$\lambda_i(t) = \lambda^{\max}\frac{S(t)}{K_i + S(t)}, \tag{S21c}$$

where: $N(t)$ is the total number of cells (per unit volume in this context); $S(t)$ is the concentration of limiting nutrient; $Y$ is the yield constant (see below); $K_i$ is the concentration of limiting nutrient that gives half the maximal growth rate constant for population $i$; $\lambda^{\max}$ is the maximal growth rate constant, assumed to be the same for all populations, for sake of simplicity; and $\mu_\lambda(t)$ is the population-mean growth rate per cell at time $t$, as defined in Eq. (S25) below.

It is worth noting that, in principle, a steady state for a chemostat with multiple populations and steady nutrient input to the chemostat is not guaranteed to exist. We show below that, if a true steady state is achieved, it is one in which only a single growth rate constant remains in the chemostat. In other words, the coexistence of multiple genotypes with different growth rate constants is not possible in a true steady state. Nevertheless, our simulations (S1.6.2) show that $N(t)$ and $S(t)$ can be very closely approximated by constants along the way towards the steady state (S1.6.2), even as population proportions evolve towards an overall steady state in which all but one growth rate constant remains in the chemostat.

The argument for the ultimate steady state, if accessible, is based on evolution equations for total cell number, $N(t)$, genotype proportions, $p_i(t)$, and the population-mean growth rate, $\mu_\lambda(t)$. (Note absence of the superscript $a$, used previously to denote arithmetic mean, as opposed to the population mean.)

As a consequence of the fact that

$$\frac{d}{dt} n_i(t) = [\lambda_i(t) - \beta] n_i(t), \tag{S22}$$

the total number of cells in the chemostat,

$$N(t) = \sum_{i=1}^{m} n_i(t), \tag{S23}$$

evolves according to

$$\frac{d}{dt} N = \left[\mu_\lambda(t) - \beta\right] N. \tag{S24}$$

In Eq (S24), $\mu_\lambda(t)$ is the population-mean fitness. It is defined by

$$\mu_\lambda(t) = \sum_{i=1}^{m} p_i(t)\, \lambda_i(t), \tag{S25}$$

where $p_i(t)$ is the proportion of cells of genotype $i$ in the chemostat at time $t$; i.e.,

$$p_i(t) = \frac{n_i(t)}{N(t)}. \tag{S26}$$

The evolution equations for $p_i(t)$, are given by taking the derivative of both sides of Eq. (S26) (using the quotient rule and Eqs. (S22)–(S25)) to obtain

$$\frac{d}{dt} p_i(t) = \left[\lambda_i(t) - \mu_\lambda(t)\right] p_i(t). \tag{S27}$$

Eq (S27) says that, at every instant, $p_i'(t)$ is proportional to $p_i(t)$ with a proportionality factor that is the deviation of fitness $\lambda_i(t)$ from the population-mean fitness, $\mu_\lambda(t)$. The implication is that, for each genotype $i$, in the steady state, where $p,'(t) = 0$ (by definition of the steady state), either $\lambda_i = \mu_\lambda$, a single number, or $p_i = 0$. Consequently, in the steady state, there is either a single $\lambda_i = \mu_\lambda$ (by definition of the mean) with corresponding $p_i = 1$, or, there are several genotypes, all with the same $\lambda_i$ value, equal to $\mu_\lambda$ (again by definition of the mean), with a pooled proportion equal to 1. In either case, a single growth rate constant remains in the steady state (if a steady state is achieved).

In our experiments with a steady input rate of a single limiting nutrient, $N(t)$ appears to be constant, long before a steady state is achieved. This is inferred from our observation of substantial changes in population proportions throughout the time course of our experiments, despite the apparent constancy of $S(t)$ and $N(t)$. What insight can theory give us about this pre-steady-state regime?

According to Eq. (S24), $N'(t) = 0$, implies that $\mu_\lambda(t) = \beta$. By the same reasoning, if $N(t)$ is approximately constant, then $\mu_\lambda(t) \approx \beta$. This approximate equality can only be maintained in the long run if $\mu'_\lambda(t) \approx 0$.

This consideration motivates a look at the evolution equation for $\mu_\lambda(t)$ to discover what it is driven by. The rate of change of $\mu_\lambda(t)$ is given by taking the derivative of both sides of Eq. (S25) (using the product rule term by term), as follows:

$$\frac{d}{dt}\,\mu_\lambda(t) = \frac{d}{dt}\sum_{i=1}^{m} p_i(t)\lambda_i(t) \tag{S28a}$$

$$= \sum_{i=1}^{m}\left[p'_i(t)\lambda_i(t) + p_i(t)\lambda'_i(t)\right] \tag{S28b}$$

$$= \left[\sum_{i=1}^{m}\left[p'_i(t)\lambda_i(t)\right]\right] + \sum_{k=1}^{m} p_k(t)\lambda'_k(t) \tag{S28c}$$

$$= \left[\sum_{i=1}^{m}\left\{\left[\lambda_i(t) - \mu_\lambda(t)\right]\, p_i(t)\right\}\lambda_i(t)\right] + \sum_{k=1}^{m} p_k(t)\lambda'_k(t) \tag{S28d}$$

$$= \left[\left\{\sum_{i=1}^{m} p_i(t)\,\lambda_i^2(t)\right\} - \mu_\lambda^2(t)\right] + \sum_{k=1}^{m} p_k(t)\lambda'_k(t) \tag{S28e}$$

$$= \sigma_\lambda^2(t) + \sum_{k=1}^{m} p_k(t)\,\lambda'_k(t). \tag{S28f}$$

Eq (S28) says that the population-mean fitness at each time $t$ changes at a rate given by the instantaneous population-variance of fitness, $\sigma_\lambda^2(t)$ plus the population-mean rate of change of the fitness. Eq (S28) is a special case of the Price equation [7] in the continuous-time form [1], in which the numerical value of the fitness trait is fitness itself. See Eq. 2.4 of [1], with the phenotypic fitness trait quantified by their variable $z$ and their net reproductive rate $r$ both equal to our fitness $\lambda$, and with no instantaneous fitness difference $\Delta z$ between parent and offspring upon birth of the offspring. Note also that the covariance term in the Price equation, $\mathrm{cov}[z, r]$ in [1], in our case, becomes $\mathrm{cov}[\lambda, \lambda] = \mathrm{var}(\lambda)$.

From Eq. (S28) one can see that $\mu'_\lambda(t)$ can be small if both terms on the right-hand side of the equation are small, or if the terms counterbalance to give a small sum. Further insight is provided below by computational modeling.

According to Eq. (S28), if a steady state does exist (where, by definition of the steady state, $\mu'_\lambda(t) = 0$, and $\lambda'_i(t) = 0$ for all $i$), in this steady state,

$$\sigma_\lambda^2 = 0. \tag{S29}$$

The variance of fitness $\lambda$ in the steady state can only be equal to zero if all the $\lambda_k$ in a set $\mathcal{K}$ for which $p_k, \neq, 0$, are equal to one and the same value — call it $\lambda_\infty$. Here, we use $\lambda_\infty$ to mean $\lim_{t\to\infty} \lambda(t)$, which could also be called $\lambda_{\rm ss}$, to refer to the steady state value. In other words, in the steady sate,

$$\lambda_k = \begin{cases} \lambda_\infty & \text{for } k \in \mathcal{K} \\ 0 & \text{otherwise} \end{cases} \tag{S30}$$

In our experimental protocol, it seems that, if there is more than one genotype with $\lambda_k = \lambda_\infty$, it is likely due to a number of gene knockouts that have no fitness phenotype.

Dean [2] discusses the practical circumstances in which multiple genotypes in a chemostat with different growth rates coexist with a fixed population number $N$. This regime corresponds to a situation in which $\mu_\lambda(t) - \beta \approx 0$, and, nevertheless, $p_i(t)$ change slowly to eventually give the true steady state in which only one growth rate constant per cell remains in the chemostat (illustrated below).

##### S1.6.2 Computational results on the possibility of fixed cell number and the coexistence of several genotypes with differing growth rate constants in the steady state

We asked about the extent to which an extension of the standard mathematical model for a chemostat [10, 2], as given in the system of equations, Eqs. (S21), can account for our experimental findings in chemostats with steady input of a single limiting nutrient.

We simulated a chemostat with 4,000 populations, each with an independent random growth rate per cell. The results we report here are generic for heterogeneous populations and not dependent on a large number of different genotypes.

We found that, following a transient period of roughy 20 generation times (Fig. S1A), $S$ and $N$ eventually settled down to nearly steady values equal to those in the true overall steady state, $S_\infty$ and $N_\infty$, respectively (Fig. S1C). Note that the $y$-axis for relative $S$ and $N$ values in Fig. S1C spans only roughly 0.99–1.125. During a 200-generation time period, after the transient, with near constancy of $S$ and $N$, the population proportions evolve slowly (Fig. S1D), and the single dominant population achieves a proportion close to 1 at the end of this time period. Fig. S1B illustrates the dramatic change in the distribution of binned population proportions between the nominal end of the transient period of 20 generation times (Fig. S1B purple bars) and end of the simulation time, 220 generation times (Fig. S1B, grey bar). At the endpoint of the simulation time, only a single gray bar is visible, and the combined proportion genotypes it comprises are accounted for almost entirely by the winning population (consistent with Fig. S1D). In Fig. S1D the blue bars reflect the distribution of random $\lambda$-values at $t = 0$, but the $\lambda_i(S)$ values are evaluated t at $S = S_\infty$. The motivation is explained below.

**Parameter choices for the simulation** were made in the following way. The "winning" population is the one with the smallest value of the half-saturating constant, which we refer to as $K_1$. This single value of $K$ was chosen deterministically. To do so we first chose the steady-state value of the limiting nutrient concentration, $S_\infty = 0.1$. We chose $K_1$ to enforce the constraint that $S_\infty = \varepsilon K_1$, where $\varepsilon << 1$; we chose $\varepsilon = 0.1$. In other words, the steady-state concentration of limiting nutrient $S_\infty$ is small compared to the half-saturating constant $K_1$. This is the typical setup of the chemostat model. We chose the single $\lambda^{\max}$ value, across populations, by satisfying the steady state condition, with only genotype 1 remaining in the chemostat. Namely that the growth rate constant is equal to the dilution rate constant; i.e., $\lambda_1(S_\infty) = \beta$. This is equivalent to

$$\lambda^{\max} \frac{S_\infty}{K_1 + S_\infty} = \beta, \tag{S31}$$

with the result that

$$\lambda^{\max} = \frac{1+\epsilon}{\epsilon}\beta. \tag{S32}$$

We chose the concentration of limiting nutrient in the input growth medium, $S_0$ to be large compared to the steady-sate value, $S_\infty$. In particular, we chose $S_0 = 10\, S_\infty$. This choice is also typical for the standard chemostat model. The steady-state density of cells in the chemostat, $N_\infty$, is determined by the yield constant $Y$ according to

$$\frac{N_\infty}{Y} = S_0 - S_\infty. \tag{S33}$$

Eq. (S33) reflects the fact that the chemostat model equations can be recast in terms of a normalized cell "density" given by $N/Y$. This means that either one can choose $N_\infty$, and this determines $Y$, or one can choose $Y$, and this determines $N_\infty$. We adopted the former approach and set $N_\infty$ to an arbitrary, but reasonable value of $2.5 \times 10^7$/mL, as in Figure 3 of [10]. The distribution of the random $\lambda_i$ values at any value of $S$ is determined, with fixed $\lambda^{\max}$, by the distribution of $K_i$ values, or vice versa. We know that the largest $\lambda(t)$, as $t \to \infty$ is

$$\lambda_1(S_\infty) = \lambda^{\max} \frac{S_\infty}{K_1 + S_\infty} = \beta. \tag{S34}$$

And we know that the other $\lambda_i(S_\infty)$ values are less than $\lambda_1(S_\infty)$. This motivates us to choose $\lambda_i(S_\infty)$ from a distribution with maximum value equal to $\beta$, and a minimum value considerably lower; we took this value to be equal to $0.1\,\beta$. We used a truncated gamma density function for sake of convenience, with a shape parameter equal to 16 (giving a standard deviation of 0.25) and a mean value equal to $\beta/2$. For any randomly chosen $\lambda_i(S_\infty)$ value, the corresponding $K_i$ value is given by

$$K_i = \left( \frac{\lambda^{\max}}{\lambda_i(S_\infty)} - 1 \right) S_\infty. \tag{S35}$$

[10] Naomi Ziv, Nathan J Brandt, and David Gresham. The use of chemostats in microbial systems biology. *J Vis Exp*, (80), Oct 2013.

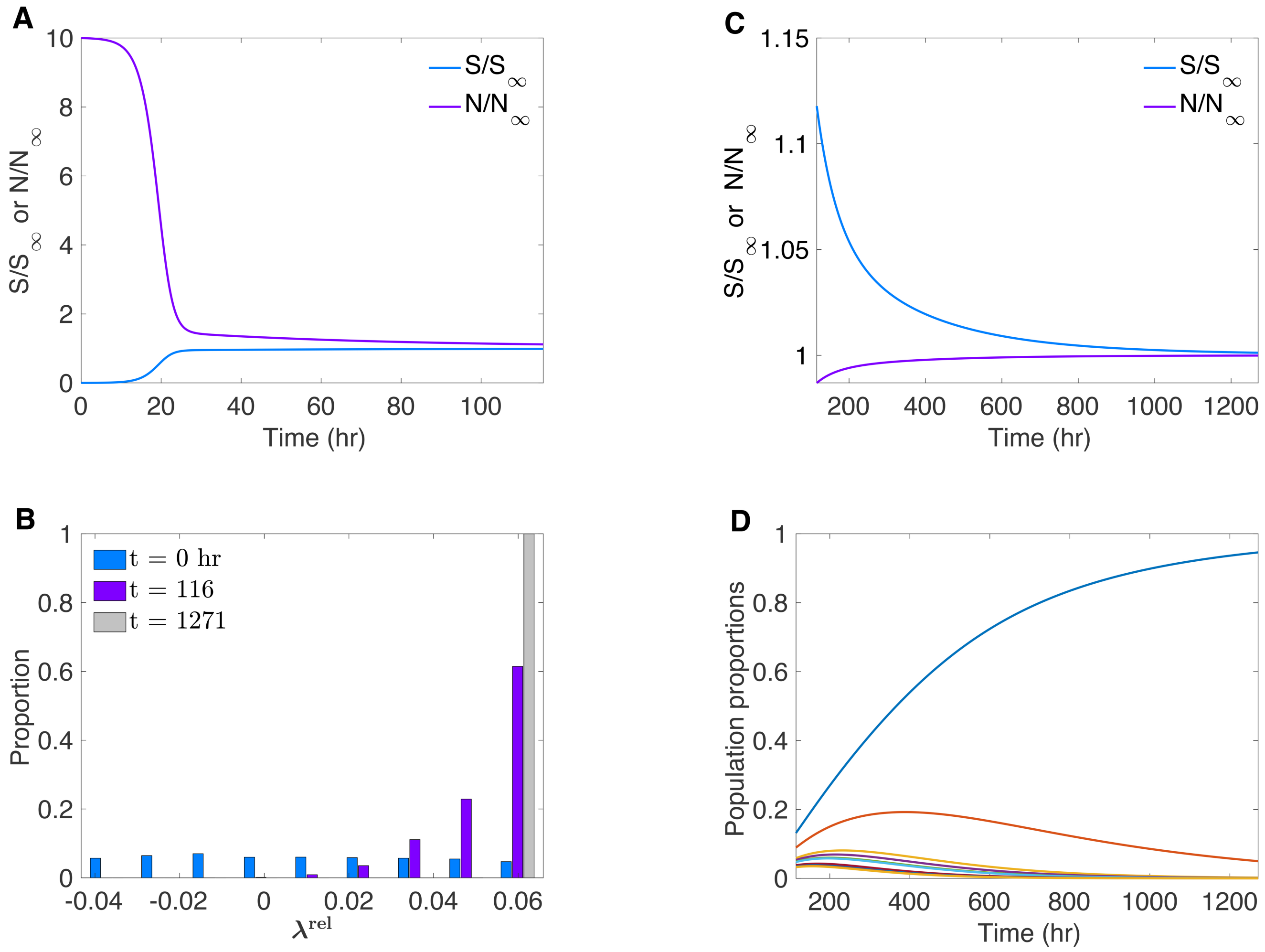


Figure S1: **Population dynamics in a chemostat with heterogeneous genotype fitness and static input of limiting-nutrient growth medium** (A) Evolution of $N(t)$ and $S(t)$ towards steady state values starting from initial conditions $N(0) << N_\infty$, and $S(0) = S_0$, the concentration of limiting nutrient in the chemostat input medium. Note that the variable are expressed in relative terms $N(t)/N_\infty$, and $S(t)/S_\infty$, so that, as $t \to \infty$, both relative values approach 1. (B) Distribution of $\lambda^{\mathrm{rel}}$ values (binned proportions) just after an initial transient time of 20 generation times (purple bars) and at 200 generation times thereafter (gray bars). Blue bars reflect distribution of $\lambda$ values at $t = 0$, as explained in the text. (C) Relative $N(t)$ and $S(t)$, showing slow approach towards a value of 1 with very small deviations over a 200-generation time period following an initial transient period. (D) Slow evolution of population proportions (top 10), while $N(t)$ and $S(t)$ hardly deviate form their steady-state values.
